# invertmeeg: A Benchmark and Unified Python Library for EEG Inverse Solvers

**DOI:** 10.64898/2026.03.06.710103

**Authors:** Lukas Hecker

## Abstract

Electroencephalography (EEG) source imaging remains difficult to compare systematically because inverse solvers are distributed across different software packages, programming languages, and evaluation protocols. We present a frozen four-scenario EEG benchmark of 106 solvers evaluated on a shared BioSemi-32 / `ico3` setup, together with `invertmeeg`, an open-source Python package that currently exposes 118 inverse solvers through a consistent two-step interface built on the `MNE-Python` ecosystem. The benchmark spans focal, multi-source, spatially extended, and low-SNR source configurations and uses earth mover’s distance (EMD) as the primary metric, with average precision (AP), mean localization error (MLE), and correlation used for complementary ranking. Across this benchmark, no single solver dominates every regime: flexible subspace and hybrid methods perform best overall, while Bayesian methods remain particularly competitive under extended-source and low-SNR conditions. The package is available via `pip install invertmeeg` and imported in Python as `invert`.

## 1 Introduction

The reconstruction of neural source activity from EEG sensor measurements is a fundamental problem in functional neuroimaging [1, 2]. Given a forward model that describes how source currents produce sensor-level signals, the inverse problem seeks to recover the underlying source distribution. Because the number of potential source locations (typically 10^3^–10^4^) far exceeds the number of sensors (typically 10^1^–10^2^), the problem is severely underdetermined and requires regularization or prior assumptions to obtain a unique solution [3].

Over the past four decades, a rich landscape of inverse methods has emerged. Minimum norm approaches [4] minimize the *L*_2_ norm of the source distribution; the LORETA family [5, 6] incorporates spatial smoothness constraints; beamformers [7] construct spatial filters that maximize signal power at a target location while suppressing interference; Bayesian methods [8, 9] cast the problem in a probabilistic framework with structured priors; sparse recovery algorithms [10] exploit the assumption that only a small number of sources are simultaneously active; and subspace scanning methods such as MUSIC [11] exploit the orthogonality between signal and noise subspaces. More recently, deep learning approaches have been proposed that learn the inverse mapping from data [12, 13].

Despite this wealth of methods, practitioners face a fragmentation problem. Implementations are distributed across different toolboxes—`MNE-Python` [14], Brainstorm [15], FieldTrip [16], SPM [17]—each supporting only a subset of methods, often with incompatible data formats and interfaces. Many published algorithms lack publicly available implementations entirely, or exist only as standalone scripts tied to specific datasets. This fragmentation impedes systematic comparison of methods under controlled conditions and makes it difficult for researchers to evaluate which method is most appropriate for their specific experimental paradigm.

The central contribution of this paper is a frozen EEG benchmark of 106 solvers across 4 evaluation scenarios that span focal dipoles, multiple focal sources, spatially extended patches, and low-SNR conditions. The benchmark uses a shared BioSemi-32 forward model, `ico3` source space, simulation pipeline, and evaluation protocol, allowing direct comparison across solver families. To make this benchmark reproducible and extensible, we package the underlying infrastructure in `invertmeeg`, an open-source Python library that currently exposes 118 inverse solvers under a single, consistent application programming interface (API). All solvers inherit from a common `BaseSolver` class and follow a two-step workflow: (1) compute an inverse operator from a forward model, and (2) apply the operator to EEG data. The package builds on `MNE-Python` for data structures, forward modeling, and visualization, ensuring compatibility with existing analysis pipelines. Source code, documentation, and tutorials are available at https://github.com/LukeTheHecker/ invertmeeg and https://lukethehecker.github.io/invertmeeg/. To provide an intuition for what these benchmark differences look like in practice, Figure 1 contrasts twelve solver families on the same extended-source EEG sample before the paper moves to aggregated rankings and scenario-level summaries.

**Figure 1:**
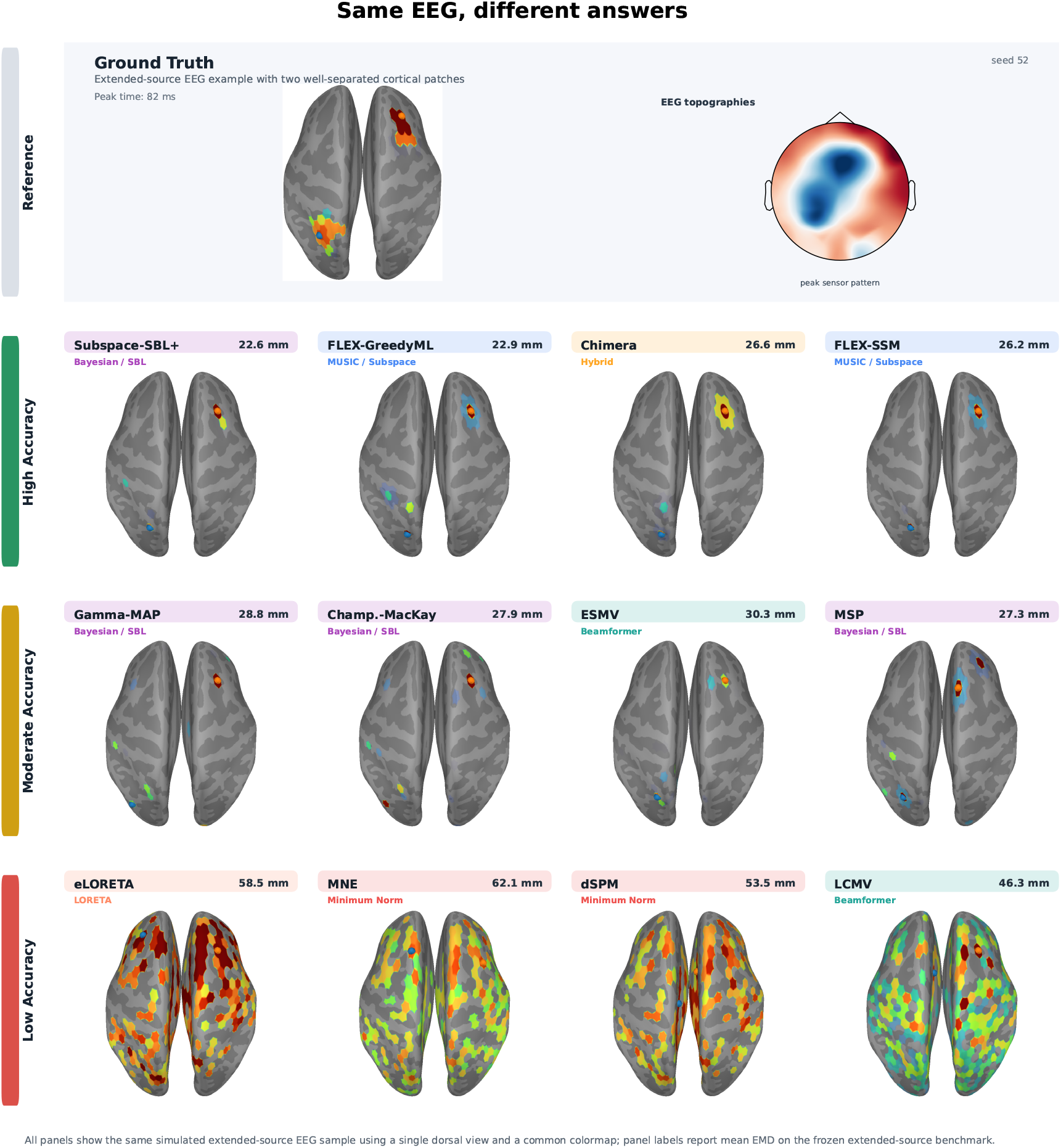
Qualitative solver spectrum for one representative extended-source EEG sample from the frozen benchmark. The ground-truth row shows two well-separated contiguous Gaussian patches together with the sensor topography at the selected peak time. The remaining panels apply twelve benchmarked solvers to the same simulated EEG data and annotate each reconstruction with its solver family and mean EMD on the frozen extended-source scenario. Row labels summarize relative performance on this specific scenario, not a universal ordering across all EEG source priors. In particular, the example favors methods that preserve compact patch structure; broader priors can be more appropriate when the underlying activation is genuinely diffuse.

Beyond the benchmark, `invertmeeg` provides:

- A**simulation framework** with diffusion-basis and contiguous Gaussian patch sources plus realistic structured sensor noise for generating diverse synthetic EEG data with known ground truth.
- **Automatic regularization** via generalized cross-validation (GCV), L-curve, and product methods, with a shared implementation across all linear solvers.
- A**standardized evaluation suite** with metrics including earth mover’s distance (EMD), average precision (AP), mean localization error (MLE), correlation, and spatial dispersion.

The remainder of this paper is organized as follows. Section 2 describes the software architecture and the solver interface. Section 3 provides an overview of the implemented solver categories. Section 4 details the shared regularization framework. Section 5 presents the simulation methodology. Section 6 describes the evaluation metrics, benchmark scenarios, and results. Section 7 provides usage examples. Section 8 discusses solver selection guidelines, related work, and limitations. Section 9 concludes the paper.

## 2 Software Architecture

A meaningful benchmark requires that all solvers are compared under identical conditions with a shared forward model, simulation pipeline, and evaluation protocol. The design of `invertmeeg` is guided by two principles: (1) every solver, regardless of its algorithmic family, should expose the same interface, and (2) adding a new solver should require minimal boilerplate. This section describes the main architectural components. Figure 2 provides an overview of the architecture and the two-step workflow.

**Figure 2:**
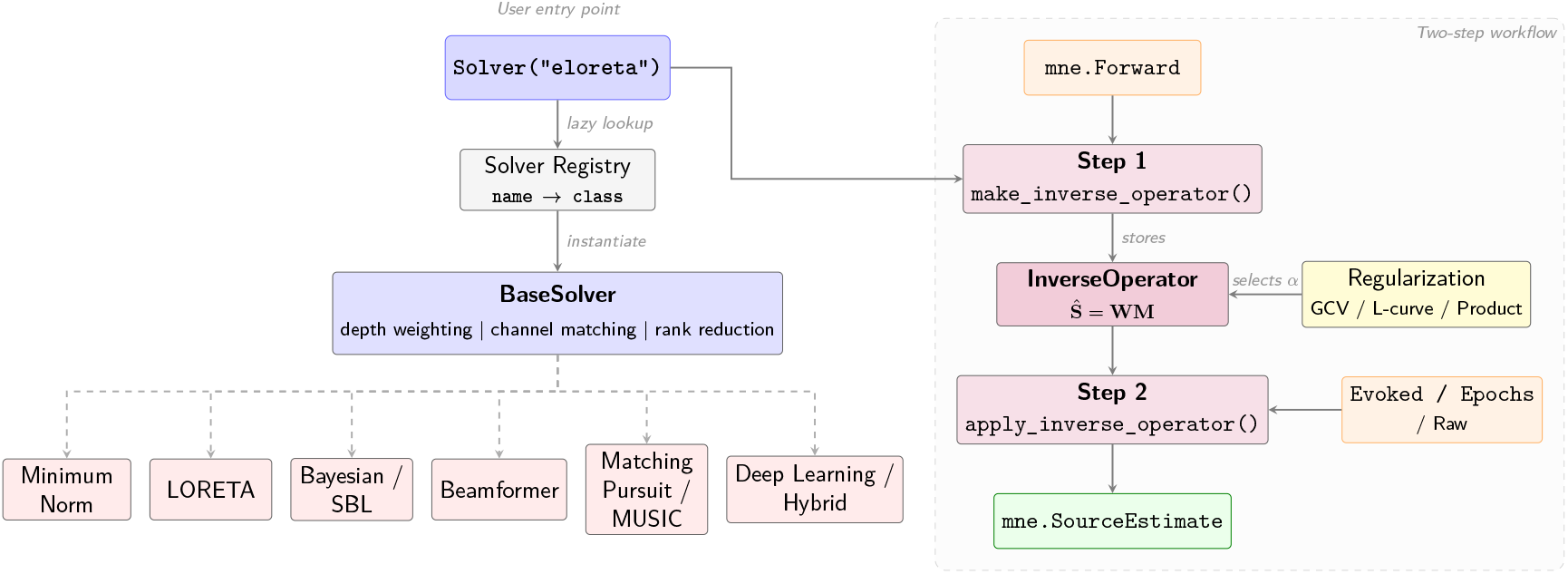
Architecture of `invertmeeg`. The `Solver()` factory lazily resolves a name to a concrete solver class inheriting from `BaseSolver`. The two-step workflow separates inverse operator computation (Step 1, depending only on the forward model) from application to data (Step 2). Regularization parameter selection happens at apply time.

### 2.1 The Solver Factory

The primary user-facing entry point is the `Solver()` factory function:

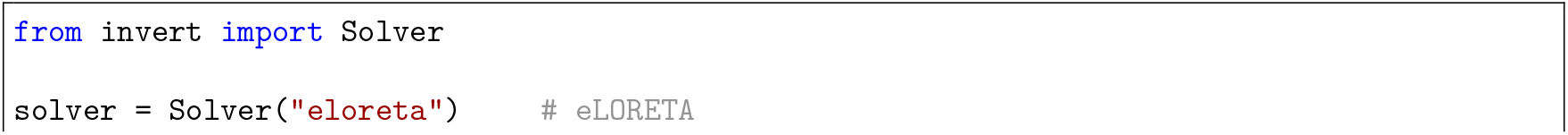
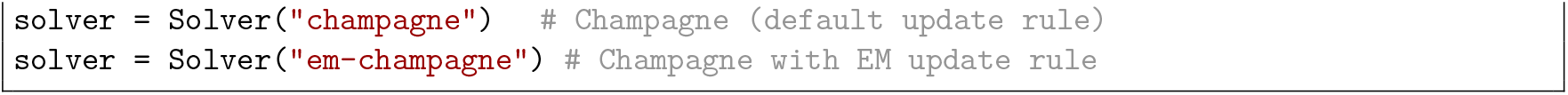

The package is installed via `pip install invertmeeg` but imported as `invert`; the code snippets below follow that Python namespace.

The factory maintains a registry that maps solver names and aliases to constructor functions. Many solvers are registered under multiple aliases for convenience (e.g., “l1”, “`fista`”, and “`mce`” all resolve to the same minimum current estimate solver). The registry is populated lazily: solver modules are imported only when the factory is first invoked, which keeps the initial import of `invertmeeg` lightweight. For solvers that represent parametric variants of the same algorithm (e.g., the seven Champagne update rules), the registry stores `functools.partial` objects that pre-fill the relevant keyword arguments.

Calling `list_solvers()` returns a sorted list of all registered aliases.

### 2.2 BaseSolver and the Two-Step Interface

Every solver class inherits from `BaseSolver`, which defines the contract:

1. `make_inverse_operator(forward, alpha=`“`auto`”, …) accepts an `MNE-Python Forward` object, computes the inverse operator, and stores it internally.
2. `apply_inverse_operator(mne_obj)` accepts an `MNE-Python` data object (`Evoked`, `Epochs`, or `Raw`), applies the precomputed inverse operator, and returns an `mne.SourceEstimate`.

This separation has a practical motivation: the inverse operator depends only on the forward model and regularization parameters, not on the data. Once computed, it can be applied to multiple recordings without recomputation, which is particularly relevant for beamformers and minimum norm methods where the operator is a matrix multiplication.

The base class handles several concerns shared across all solvers:

- **Forward model preparation**: conversion of free-orientation forward models to fixed orientation, optional common average referencing of the leadfield, and depth weighting of leadfield columns.
- **Channel matching**: automatic intersection of channels present in the forward model and the data object.
- **Regularization parameter grid**: construction of a logarithmically spaced grid of regularization parameters, scaled by the largest eigenvalue of the leadfield Gram matrix **LL**^⊤^ (see Section 4).
- **Rank reduction**: optional signal subspace selection via SVD-based rank estimation.
- **Output conversion**: mapping of the source matrix back to an `mne.SourceEstimate` with correct vertex indices, timing, and subject metadata.

### 2.3 InverseOperator

For solvers that produce a linear inverse operator (a matrix 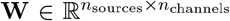), the precomputed operator is wrapped in an `InverseOperator` object. Applying the operator to a data matrix 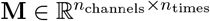 is then a single matrix multiplication:

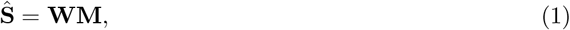

where 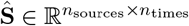 is the estimated source activity. When automatic regularization is enabled (`alpha=`“`auto`”), the solver stores one `InverseOperator` per candidate regularization parameter; the optimal one is selected at apply time based on the chosen regularization criterion (Section 4).

### 2.4 Solver Metadata

Each solver class carries a `SolverMeta` dataclass with fields for the full solver name, acronym, category, a brief description, and literature references. This metadata serves two purposes: it enables programmatic generation of solver catalogs and documentation, and it allows users to inspect a solver’s provenance at runtime.

### 2.5 Depth Weighting

Sources deeper in the brain produce weaker signals at the sensor level, biasing unweighted solutions toward superficial cortex. `invertmeeg` applies column-wise depth weighting to the leadfield matrix **L** before computing the inverse operator:

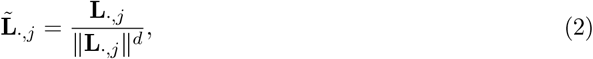

where *d* ∈ [0, 1] controls the weighting strength (default *d* = 0.5). After source estimation, the weighting is removed to recover physical amplitude units.

## 3 Solver Catalog

`invertmeeg` currently registers 118 solver implementations. The frozen EEG benchmark reported here evaluates 106 of them under a common four-scenario setup. Rather than presenting the full mathematical derivation of each solver, we summarize the main families and refer the reader to the original publications. A complete solver reference with canonical identifiers and aliases is provided in the Appendix and as a machine-readable supplementary CSV.

### 3.1 Minimum Norm Methods

Minimum norm methods seek the source distribution ŝ that minimizes a weighted norm of the source vector subject to the data fit:

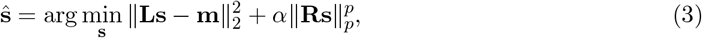

where **L** is the leadfield matrix, **m** is the measurement vector, *α* is the regularization parameter, **R** is a weighting matrix, and *p* ∈ {1, 2} selects the norm.

For *p* = 2 and **R** = **I**, Equation (3) yields the classic minimum norm estimate (MNE) [4] with the closed-form solution:

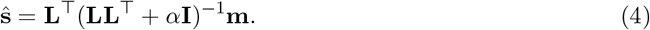

Variants include weighted MNE (wMNE), which uses noise-weighted columns, and dynamic statistical parametric mapping (dSPM) [18], which normalizes source power by noise variance.

For *p* = 1, Equation (3) promotes sparse solutions; `invertmeeg` implements this via the FISTA algorithm [19] as well as combined *L*_1_*/L*_2_ penalties.

### 3.2 LORETA Family

Low Resolution Electromagnetic Tomography (LORETA) [5] adds a spatial smoothness constraint by incorporating the cortical graph Laplacian **Δ** into the regularization term. The family includes LORETA, standardized LORETA (sLORETA) [6], and exact LORETA (eLORETA) [20], which differ in their normalization strategies and have distinct localization properties.

### 3.3 Bayesian Methods

Bayesian approaches model the source distribution as a random variable with a structured prior and optimize the model evidence (type-II maximum likelihood). The Champagne algorithm [8] and its variants form the largest subgroup, with seven update rules (MacKay, Convexity/MM, EM, AR-EM, TEM, LowSNR, Adaptive) and optional noise learning. Other Bayesian solvers include Gamma-MAP [21], Multiple Sparse Priors (MSP) [9], coherent maximum entropy on the mean (cMEM) [22], and Sparse Bayesian Learning (SBL) variants.

### 3.4 Beamformers

Beamformers construct spatial filters that pass signals from a target location while minimizing contributions from elsewhere. The linearly constrained minimum variance (LCMV) beamformer [7] solves:

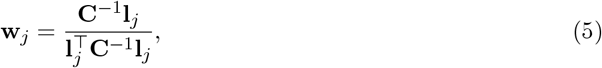

where **C** is the data covariance matrix and **l**_*j*_ is the leadfield column for source *j*. `invertmeeg` implements 25 beamformer variants, including DICS for frequency-domain analysis [23], synthetic aperture magnetometry (SAM) [24], eigenspace-projected variants (ESMV family), multiple-constraint approaches (MCMV), and the empirical Bayesian beamformer (EBB).

### 3.5 Matching Pursuit and Sparse Recovery

These methods greedily or iteratively select active sources from the leadfield dictionary. Orthogonal matching pursuit (OMP) [25] iteratively selects the leadfield column most correlated with the residual; CoSaMP [26] and subspace pursuit [27] extend this with two-stage selection and pruning. `invertmeeg` includes 10 matching pursuit variants, including FLEX variants that combine greedy selection with flexible subspace scanning, ReMBo (random embedding), and hierarchical subspace methods (SubSMP, ISubSMP).

### 3.6 MUSIC and Subspace Methods

Multiple signal classification (MUSIC) [11] localizes sources by scanning for leadfield columns that lie in the signal subspace. RAP-MUSIC [28] extends this with recursive projection to resolve multiple sources. `invertmeeg` provides 13 subspace methods: MUSIC, RAP-MUSIC, TRAP-MUSIC, and their FLEX variants, as well as signal subspace matching (SSM), alternating projections (AP), ExSo-MUSIC, and maximum-likelihood scanners (ExhaustiveML, GreedyML) that search for the source configuration maximizing the data likelihood.

### 3.7 Dipole Fitting

Equivalent current dipole (ECD) fitting models the source as one or more point dipoles whose positions and moments are optimized to fit the data. SESAME [29] uses a sequential Monte Carlo approach for multi-dipole estimation.

### 3.8 Deep Learning

`invertmeeg` includes six neural network architectures that learn the inverse mapping from simulated training data. These solvers require PyTorch as an optional dependency and use the simulation framework (Section 5) for on-the-fly training data generation—no pre-recorded data is needed. Three of these solvers (FC, CovCNN, CovCNN-KL) are included in the benchmark.

**FC** is a simple fully connected EEG inverse-model baseline in the ConvDip / ESINet family of early deep-learning approaches [12, 13]. Each time point is processed independently through a dense layer (*n*_channels_ → *n*_units_, tanh activation), a second dense layer (*n*_units_ → *n*_units_), and a linear output layer (*n*_units_ → *n*_dipoles_). With 600 hidden units and 1,284 dipoles, this yields ∼1.2 × 10^6^ trainable parameters.

**CovCNN** [13] operates on the sensor covariance matrix 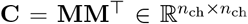 rather than the raw time series, providing implicit temporal averaging and sign invariance. The covariance is passed through a frozen convolutional layer whose weights are derived from the leadfield matrix, projecting the *n*_ch_ × *n*_ch_ input to *n*_dipoles_ feature maps. The resulting features are flattened and processed by a multi-layer perceptron (MLP) with tanh activations, producing per-dipole source estimates. CovCNN is trained with cosine similarity loss.

**CovCNN-KL** shares the CovCNN architecture but is trained with Kullback-Leibler divergence loss after applying a softmax to the output logits, treating source localization as a distribution-matching problem. This reformulation improves localization accuracy compared to the regression-based CovCNN. In the benchmark configuration (24 hidden units), both CovCNN variants have ∼1.0 × 10^6^ trainable parameters; FC has ∼1.2 × 10^6^. All three are trained with the Adam optimizer for 200 epochs with early stopping (patience 60) and a batch size of 2,048 simulated samples.

The remaining (non-benchmarked) architectures include CovCNN-KL-FlexOMP, which augments CovCNN-KL with iterative sparse post-processing; a bidirectional LSTM for temporal modeling; and a CNN with LSTM encoder for spatiotemporal processing.

### 3.9 Flexible Extent Estimation (FLEX)

A common limitation of classical inverse methods is the assumption that each source is a point dipole. In practice, cortical generators often extend over patches of varying size. The FLEX (Flexible Extent Estimation) framework [30] addresses this by augmenting any base solver with a multi-order leadfield dictionary that allows simultaneous estimation of source location and spatial extent.

The core idea is to construct smoothed versions of the leadfield matrix **L** via iterative diffusion on the cortical adjacency graph. Given the graph Laplacian **Δ** of the source space, a smoothing operator is defined as:

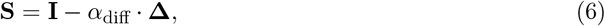

where *α*_diff_ controls the diffusion rate. Applying **S** to the leadfield produces leadfields for increasingly extended sources:

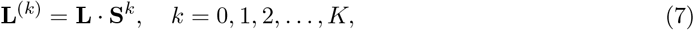

where *k* = 0 corresponds to point dipoles and increasing *k* produces progressively smoother spatial profiles. For each order *k*, a gradient matrix **B**^(*k*)^ tracks how smoothed activations map back to individual dipoles in the original source space.

The extended leadfield dictionary **L**_ext_ = [**L**^(0)^ | **L**^(1)^ | · · · | **L**^(*K*)^] contains *K* · *n* candidate columns (one per dipole per order). The base solver then operates on this extended dictionary and selects the best (location, order) pair for each source using its native selection criterion—subspace projection for MUSIC variants, residual correlation for matching pursuit, likelihood maximization for ML scanners, evidence maximization for Bayesian methods, or beamformer output power for spatial filters. A mutual exclusivity constraint ensures that at most one order is active per dipole location. The selected sources are mapped back to the original source space via their corresponding gradient matrices **B**^(*k*)^.

This mechanism is general: `invertmeeg` applies it to subspace methods (FLEX-MUSIC, FLEX-SSM, FLEX-GreedyML), matching pursuit algorithms (FLEX-OMP, FLEX-CoSaMP, FLEX-SP, FLEX-ReMBo), beamformers (the FLEX-ESMV family), and Bayesian solvers (Flex-Champagne, Flex-NL-Champagne). In all cases, the data-driven order selection allows each source to adopt the spatial extent that best explains the measurements, without requiring the user to specify source size a priori. Across the frozen EEG benchmark, several FLEX variants rank near the top overall, particularly in the focal and multi-source regimes.

### 3.10 Hybrid Methods

`invertmeeg` includes three hybrid solvers that combine multiple algorithmic strategies to adapt to the characteristics of the data at hand.

**Chimera** adaptively selects between a beamformer (FLEX-ESMV) and a subspace scanner (SSM) based on properties estimated from the data. It first estimates the number of active sources 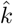 using the minimum description length (MDL) criterion [31] applied to the eigenvalues of the data covariance. If exactly one source is detected 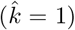, Chimera computes an order-score ratio that compares the subspace fit of a point-dipole leadfield against smoothed (extended) leadfields: if the ratio indicates a focal source, the beamformer branch is used; otherwise, SSM handles multi-source or extended configurations. This two-stage decision rule allows Chimera to route each data sample to the solver best suited to its source geometry.

**Hydra** combines a flexible maximum-likelihood scanner (FLEX-GreedyML) with an adaptive beamformer (Adapt-FLEX-ESMV), switching between them based on a variance-based heuristic. The mean sensor variance of the input data is compared against a calibrated threshold: high-variance data (characteristic of distributed or ongoing activity) is routed to the beamformer, while lower-variance data (characteristic of sparse focal sources) is processed by FLEX-GreedyML. In the frozen EEG benchmark snapshot, Hydra ties for the best global rank, illustrating how a simple switching heuristic can leverage the complementary strengths of sparse maximum-likelihood scanning and adaptive beamforming.

**APSE** (Alternating Projections with Source Enumeration) integrates subspace scanning with automatic model-order selection via information-theoretic criteria.

### 3.11 Novel Solvers Developed with AI Coding Agents

Beyond re-implementing published algorithms, `invertmeeg` includes more than 30 currently unpublished solver variants. These methods were developed by the first author in collaboration with AI coding agents (large language models used as interactive programming assistants) and represent novel combinations and extensions of existing algorithmic components. Families of novel solvers include FLEX-ESMV beamformer variants, Flex-Champagne Bayesian solvers, hybrid switching methods (Chimera, Hydra, APSE), subspace matching pursuit variants, and all deep learning solvers (CovCNN, CovCNN-KL, CNN, LSTM). Each novel solver is documented with its component references in the solver source code and in the appendix tables.

As an illustrative example, we describe **Subspace-SBL**—one of the strongest Bayesian solvers and a top-five method in the frozen benchmark—in detail. Subspace-SBL decomposes the inverse problem into two stages: a combinatorial source detection stage followed by a continuous amplitude estimation stage on the detected support.

#### Stage 1: Subspace detection

Given sensor data **Y** ∈ ℝ^*m*×*T*^, we first form a regularized signal-subspace projector via signal subspace matching (SSM):

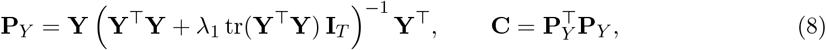

where *λ*_1_ is a small regularization constant. The matrix **C** captures the signal subspace robustly even under model errors. Sources are then detected greedily from the multi-order FLEX dictionary {**L**^(0)^, …, **L**^(*K*)^}. At each step *q*, we select the (location, order) pair that maximizes the projected subspace score:

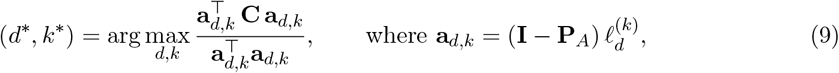

and **P**_*A*_ is the (regularized) projector onto the subspace spanned by previously selected leadfield columns. After the greedy forward pass, a cyclic refinement procedure revisits each selected source and re-optimizes its (location, order) assignment while holding the others fixed, iterating until convergence.

#### Stage 2: Noise-learning SBL refinement

Let 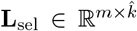 denote the leadfield columns corresponding to the 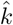 detected sources. We model the data as **Y** = **L**_sel_**S** + **N**, with a Gaussian prior **s**_*t*_ ∼ 𝒩 (**0**, diag(*γ*)) and heteroscedastic sensor noise **n**_*t*_ ∼ 𝒩 (**0**, diag(*λ*)). The model covariance is 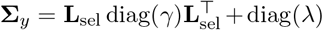. Source variance and noise parameters are estimated by iterating Type-II ML fixed-point updates:

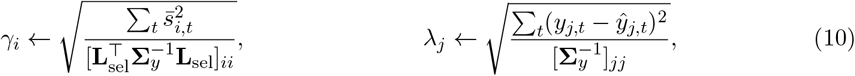

where 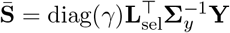 is the posterior mean and 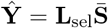 the predicted data. Sources with *γ*_*i*_ < 10^−3^ max(*γ*) are pruned, and the algorithm converges when the relative change in the Type-II ML objective falls below 10^−8^. The final inverse operator is 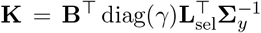, where **B** contains the FLEX gradient vectors that map detected atoms back to the full source space.

**Subspace-SBL+** extends this by expanding the SSM-detected candidate set with graph-neighborhood atoms (one hop in the cortical adjacency graph) before running the SBL refinement. This local expansion addresses SSM’s sensitivity to vertex-level misalignment: the Bayesian refinement automatically prunes spurious candidates while retaining those that genuinely explain data variance.

The design philosophy behind these variants follows a common pattern: combine fast geometric methods (subspace scanning, beamforming) for discrete decisions with statistically regularized refinement (SBL or maximum likelihood) for continuous estimation. A dedicated paper on the agentic development process is in preparation.

## 4 Regularization

Many inverse solvers require a regularization parameter *α* that balances data fit against solution complexity. Choosing *α* appropriately is critical: too little regularization amplifies noise, while too much regularization oversmooths the solution. `invertmeeg` provides a shared regularization framework that is used by all solvers that produce linear inverse operators.

### 4.1 Regularization Parameter Grid

When `alpha=`“`auto`” (the default), the solver constructs a grid of candidate values. A set of dimensionless knobs *r*_*i*_ is logarithmically spaced in [10^−10^, 10^1^], and each is scaled by the largest eigenvalue *λ*_max_ of a reference matrix (by default **LL**^⊤^):

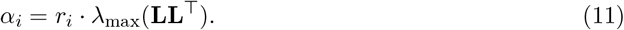

This scaling ensures that the regularization grid is adapted to the magnitude of the forward model, making the same *r*-values meaningful across different sensor configurations and source spaces. For each *α*_*i*_, a separate inverse operator **W**_*i*_ is precomputed and stored.

### 4.2 Generalized Cross-Validation (GCV)

GCV [32] selects the *α* that minimizes a leave-one-out prediction error estimate without explicit knowledge of the noise level. For a linear inverse operator that produces the hat matrix **H**_*α*_ = **LW**_*α*_, the GCV criterion is:

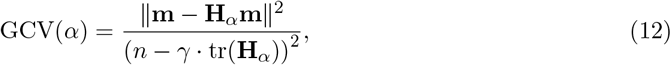

where *n* is the number of channels and *γ* ≥ 1 is a correction factor. Setting *γ* = 1 gives standard GCV; values slightly above 1 (modified GCV, MGCV) bias the selection toward stronger regularization, which can be helpful when the forward model is rank-deficient.

For iterative Bayesian solvers such as MacKay-Champagne and Convexity-Champagne, standard GCV selected unstable regularization values in our benchmark setup and led to degraded localization quality. To address this, we use the strong robust GCV (R1GCV) variant [33]:

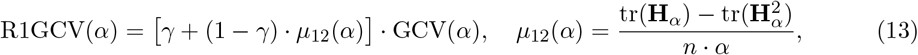

where *µ*_12_ is the negative derivative of the mean leverage tr(**H**_*α*_)/*n* with respect to *α*. The weighting factor penalizes regions of the regularization grid where solution variance amplifies rapidly as *α* decreases, biasing the selection toward more stable operating points. We therefore use R1GCV as an empirical stability choice for the Champagne-family benchmark runs and do not claim a more general mechanistic explanation here.

### 4.3 L-Curve Method

The L-curve method [34] plots the solution norm ∥ŝ∥_2_ against the residual norm ∥**L**ŝ − **m**∥_2_ for each candidate *α*. The optimal *α* is located at the “corner” of the resulting L-shaped curve, which represents the best trade-off between data fit and solution regularity. `invertmeeg` detects the corner using a triangle-area heuristic: the candidate that maximizes the area of the triangle formed with the two endpoints of the curve is selected.

### 4.4 Product Method

The product method [3] selects the *α* that minimizes the product of the solution norm and the residual norm:

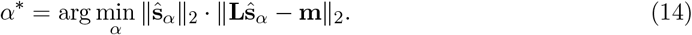

This criterion does not require computing the hat matrix and is therefore computationally cheaper than GCV.

## 5 Simulation Methodology

A meaningful comparison of inverse solvers requires synthetic data with known ground truth that spans the diversity of source configurations encountered in practice. Existing simulation approaches often rely on simple Gaussian patches centered on fixed cortical locations, which produces unrealistically regular spatial patterns and may favor solvers whose assumptions match these idealized geometries. We designed a simulation framework that generates more diverse and challenging source configurations through two complementary spatial models, structured sensor noise, and flexible temporal dynamics.

### 5.1 Spatial Source Models

`invertmeeg` provides two spatial source models that produce qualitatively different source geometries.

#### Diffusion basis patches

The first model constructs source patterns using a graph-based spatial basis derived from the cortical mesh adjacency. Given the adjacency matrix **A** of the source space, the diffusion-based gradient operator is:

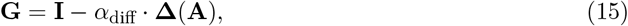

where **Δ**(**A**) is the graph Laplacian and *α*_diff_ controls the smoothing strength. Multi-order spatial patterns are generated by iteratively applying **G** to a seed dipole, where the spatial order parameter *k* controls source extent: *k* = 0 produces point dipoles, *k* = 1−2 produces cortical patches, and *k* ≥ 3 produces broadly distributed activity.

#### Contiguous Gaussian patches

The second model generates spatially extended sources with irregular, anatomically plausible geometries. For each source, a contiguous region of a specified size is grown from a random seed vertex using randomized breadth-first search (BFS) on the cortical adjacency graph. At each step, a random vertex is selected from the frontier (the set of unvisited neighbors of the current region), producing irregular region shapes that conform to the cortical surface topology rather than forming symmetric discs.

Within each region, one or two spatial components are created with Gaussian-weighted profiles. For each component, a center vertex *c* is chosen and the graph hop distance *d*_*v*_ from *c* to every vertex *v* in the region is computed via BFS restricted to the region. The spatial weights follow a Gaussian kernel:

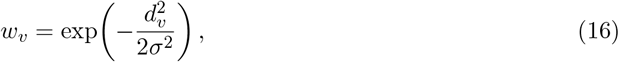

where *σ* controls the spatial smoothness (in graph-hop units). For rank-2 patches, a second component with a different center produces two overlapping smooth maps within the same region, yielding more complex spatial patterns.

This approach produces sources that are (1) contiguous and connected on the cortical surface, (2) irregularly shaped due to the randomized BFS growth, and (3) smoothly varying within the patch rather than sharply bounded. These properties make contiguous Gaussian patches substantially more diverse than standard Gaussian blobs and more representative of the spatially extended generators observed in electrophysiological recordings of oscillatory or evoked activity.

### 5.2 Temporal Dynamics

Source time courses are drawn from a pre-generated pool of colored noise signals with power spectral density *S*(*f*) ∝ 1/*f* ^*β*^, where *β* is uniformly sampled from a configured range. This produces time courses ranging from white noise (*β* = 0) to smooth, slowly varying signals (*β* = 3). Inter-source correlations are introduced via Cholesky decomposition of a correlation matrix, allowing control over the degree of temporal dependence between simultaneously active sources.

### 5.3 Sensor Noise Model

Sensor-level noise is added at a specified signal-to-noise ratio (SNR in dB). For the benchmark evaluation, we use a “realistic” noise model that combines three components to approximate the structure of empirical EEG noise:

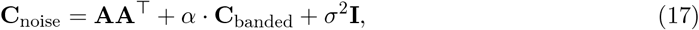

where **A** ∈ ℝ^*m*×*r*^ is a random basis generating *r* low-rank artifact-like components (simulating environmental interference or physiological artifacts), **C**_banded_ captures spatial correlations between neighboring channels (with elements [**C**_banded_]_*ij*_ = *ρ*^|*i*−*j*|^ for a coefficient *ρ*), and *σ*^2^**I** adds a white noise floor. All components are trace-normalized before combination to keep their relative contributions interpretable. The low-rank dimension *r* is randomly drawn from a configured range (typically 2−8), introducing variability in the noise structure across simulations.

This structured noise model is more challenging than i.i.d. Gaussian noise because it introduces spatial correlations and low-rank interference patterns that can bias inverse solutions toward artifactual source configurations, particularly for methods that assume spatially white noise.

### 5.4 Simulation Modes

Source activity in the benchmark consists of the selected source patches, representing the common assumption that a small number of discrete cortical generators are active. The framework additionally supports a mixture mode that combines sparse patches with smooth spatiotemporal background activity, but this mode is not used in the current benchmark.

## 6 Evaluation and Benchmarking

`invertmeeg` provides a standardized evaluation suite for comparing solver performance on simulated data with known ground truth.

### 6.1 Metrics

The following metrics are computed between the true source estimate **y**_true_(*t*) and the predicted estimate **y**_pred_(*t*). For the localization-style metrics (EMD, AP, MLE, and SD), the implementation first collapses each spatiotemporal estimate to a single unsigned spatial map by averaging absolute activity across time,

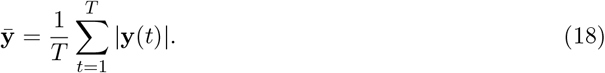

Distances are computed from Euclidean distances between dipole positions and reported in millimeters in the tables and figures. Correlation is the only metric that operates on the full time series. We designate EMD as the primary metric and AP as the secondary metric for ranking solvers, as they capture complementary aspects of reconstruction quality.

#### Earth Mover’s Distance (EMD, primary metric)

We compute the Wasserstein-1 distance between the collapsed maps 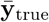 and 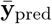 using optimal transport [35]. Before transport, values below 25% of the maximum of each map are set to zero, and the remaining mass is normalized to sum to one. This suppresses low-amplitude diffuse tails that would otherwise dominate transport cost despite contributing little to the main reconstruction. The EMD then quantifies the minimum spatial “work” required to transform one thresholded source distribution into the other, jointly capturing localization error, missed mass, false positives, and extent mismatch.

#### Average Precision (AP, secondary metric)

Average precision summarizes how well a solver ranks truly active dipoles above inactive ones across all score thresholds. The ground-truth support is binarized from 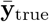 using a relative threshold of 10% of its maximum. The prediction scores are the entries of 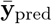. Dipoles are sorted by predicted score, a precision-recall curve is traced from that ordering, and AP is computed as the area under that curve. We prefer AP over the more commonly reported area under the ROC curve (AUC) because source imaging is an inherently imbalanced classification problem: in a source space with *n* = 1,284 dipoles, a focal source activates only 1 dipole (< 0.1% positive rate), and even extended sources rarely exceed 3–5% of the source space. Under such extreme class imbalance, AUC is dominated by the true negative rate and can assign high scores to reconstructions that spread activity broadly, whereas AP focuses exclusively on the precision among predicted positives and is therefore more sensitive to false activations [36].

#### Mean Localization Error (MLE)

MLE is computed from graph-local maxima rather than from centers of mass. Local maxima are detected separately in 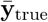 and 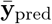 after thresholding each map at 10% of its maximum and comparing every candidate dipole against its neighbors in the cortical adjacency graph. At most five maxima are retained per map. The benchmark uses the default “DLE” aggregation mode implemented in the package: if *D*_*ij*_ is the Euclidean distance between true maximum *i* and predicted maximum *j*, then

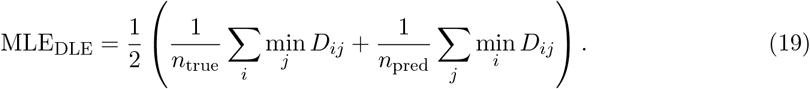

This symmetric nearest-peak distance penalizes both missed true generators and spurious predicted ones.

#### Correlation

Correlation measures temporal fidelity on the original source time series rather than on the collapsed spatial maps. For multi-time-point data, a dipole is treated as active if its maximum absolute ground-truth amplitude reaches at least 10% of the global peak over all dipoles and times. For each active dipole, we compute the Pearson correlation between its true and predicted time courses and then average across active dipoles. By default the implementation uses *unsigned* correlation, i.e. it takes the absolute value of each Pearson coefficient before averaging, so sign-flipped but otherwise correct time courses are not penalized. For single-time-point data, the code falls back to a spatial Pearson correlation across dipoles.

#### Spatial Dispersion (SD)

Spatial dispersion is reported as a blurring ratio based on a full-width-at-half-maximum-style proxy. For the collapsed true and predicted maps, nodes above 50% of the peak amplitude are identified, and a local extent measure is formed by summing distances to adjacent active nodes. SD is then the ratio of predicted extent to true extent, so values above 1 indicate a reconstruction that is spatially broader than the ground truth. We report SD descriptively but exclude it from the ranking procedure below.

### 6.2 Evaluation Scenarios

Different inverse methods are suited to different source configurations. We define four evaluation scenarios that span the range of conditions encountered in practice (Table 3):

**Table 1:** Benchmark depth-weighting policy. Category defaults apply unless a solver-specific override is listed. Solvers not shown use their package defaults without an explicit benchmark override.

| Policy | Depth weight $d$ |
| --- | --- |
| Minimum Norm default | 0.5 |
| LORETA default | 0.0 |
| Bayesian / SBL default | 0.0 |
| MUSIC / Subspace default | 0.5 |
| Backus-Gilbert override | 0.0 |
| GBF override | 1.0 |
| GFT-L1 override | 0.0 |
| MNE override | 1.0 |
| S-MAP override | 0.0 |
| TV override | 0.0 |

**Table 2:**
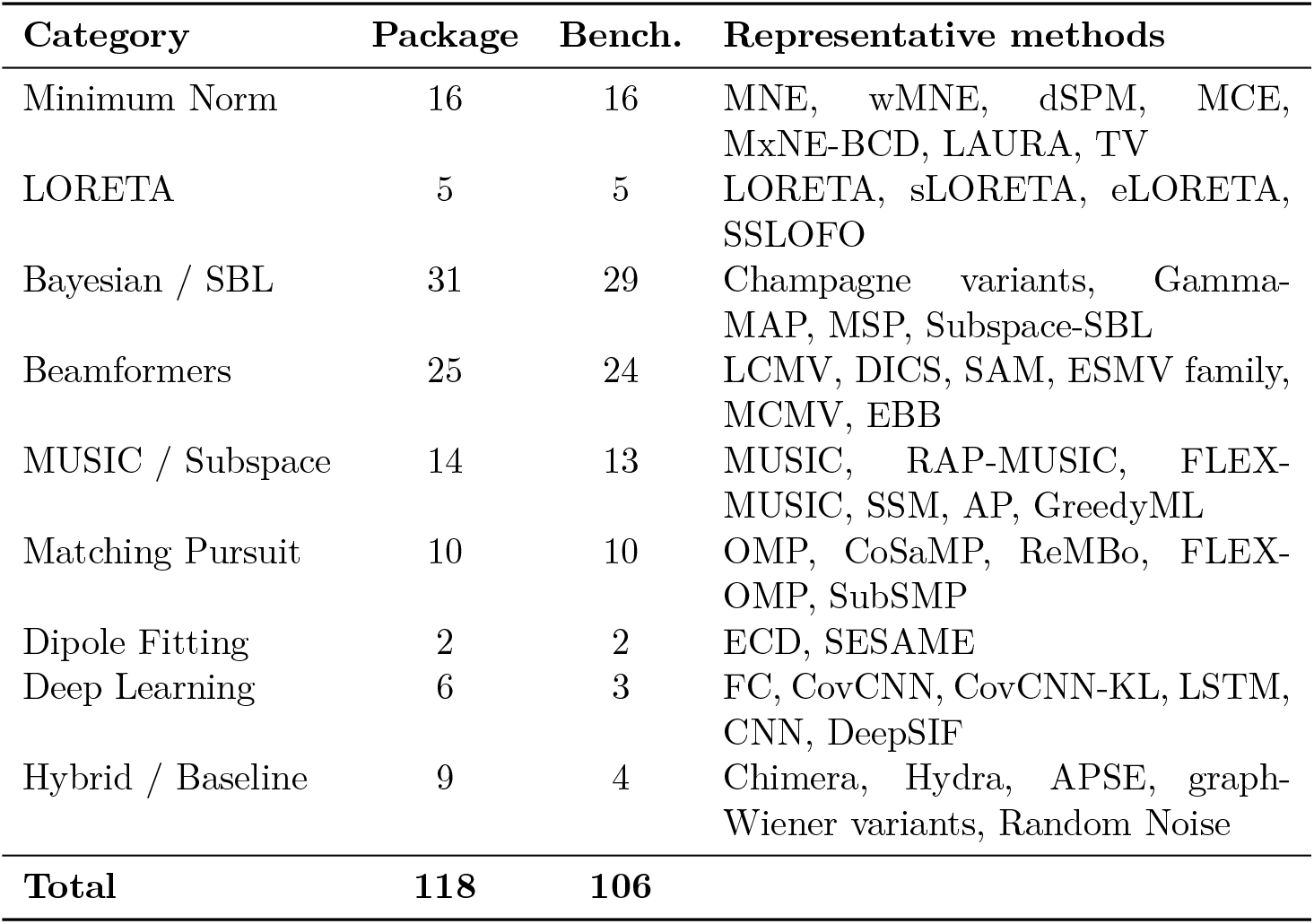
Solver categories in invertmeeg. Package totals are derived from the live registry; benchmark counts come from the frozen four-scenario EEG snapshot.

| Category | Package | Bench. | Representative methods |
| --- | --- | --- | --- |
| Minimum Norm | 16 | 16 | MNE, wMNE, dSPM, MCE, MxNE-BCD, LAURA, TV |
| LORETA | 5 | 5 | LORETA, sLORETA, eLORETA, SSLOFO |
| Bayesian / SBL | 31 | 29 | Champagne variants, Gamma-MAP, MSP, Subspace-SBL |
| Beamformers | 25 | 24 | LCMV, DICS, SAM, ESMV family, MCMV, EBB |
| MUSIC / Subspace | 14 | 13 | MUSIC, RAP-MUSIC, FLEX-MUSIC, SSM, AP, GreedyML |
| Matching Pursuit | 10 | 10 | OMP, CoSaMP, ReMBo, FLEX-OMP, SubSMP |
| Dipole Fitting | 2 | 2 | ECD, SESAME |
| Deep Learning | 6 | 3 | FC, CovCNN, CovCNN-KL, LSTM, CNN, DeepSIF |
| Hybrid / Baseline | 9 | 4 | Chimera, Hydra, APSE, graph-Wiener variants, Random Noise |
| **Total** | **118** | **106** |  |

**Table 3:**
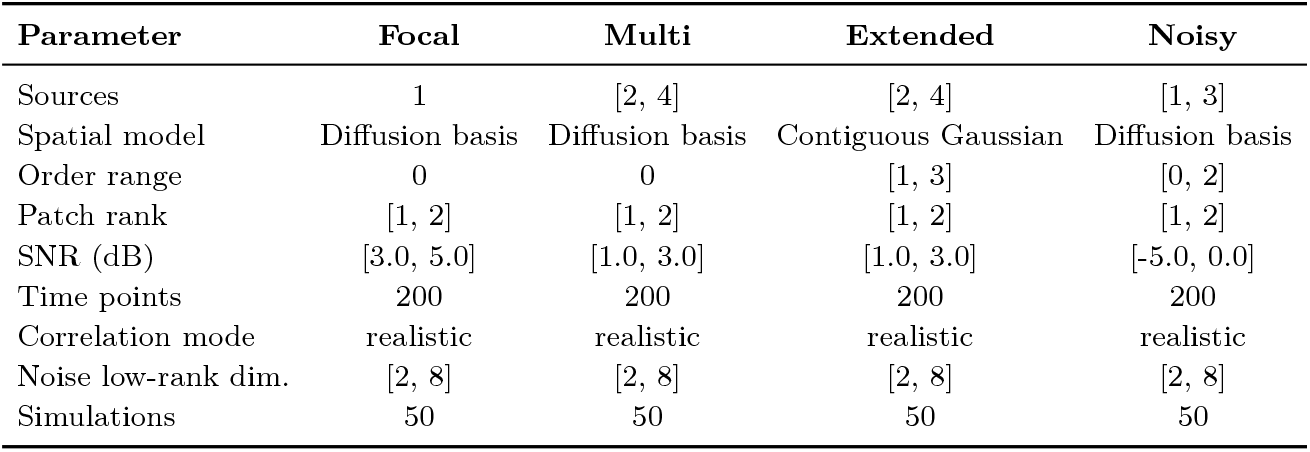
Configuration of the four EEG benchmark scenarios in the frozen paper snapshot.

| Parameter | Focal | Multi | Extended | Noisy |
| --- | --- | --- | --- | --- |
| Sources | 1 | [2, 4] | [2, 4] | [1, 3] |
| Spatial model | Diffusion basis | Diffusion basis | Contiguous Gaussian | Diffusion basis |
| Order range | 0 | 0 | [1, 3] | [0, 2] |
| Patch rank | [1, 2] | [1, 2] | [1, 2] | [1, 2] |
| SNR (dB) | [3.0, 5.0] | [1.0, 3.0] | [1.0, 3.0] | [-5.0, 0.0] |
| Time points | 200 | 200 | 200 | 200 |
| Correlation mode | realistic | realistic | realistic | realistic |
| Noise low-rank dim. | [2, 8] | [2, 8] | [2, 8] | [2, 8] |
| Simulations | 50 | 50 | 50 | 50 |

**Table 4:**
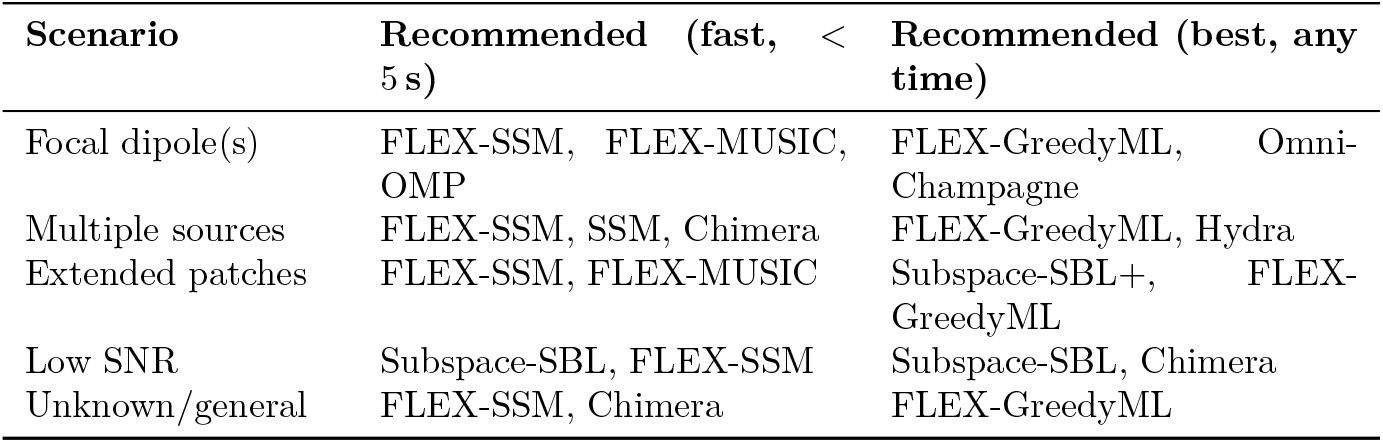
Solver selection guide based on expected source configuration and computational budget. Inference times on a single CPU core (BioSemi-32, ico3 source space).

| Scenario | Recommended (fast, < 5 s) | Recommended (best, any time) |
| --- | --- | --- |
| Focal dipole(s) | FLEX-SSM, FLEX-MUSIC, OMP | FLEX-GreedyML, Omni-Champagne |
| Multiple sources | FLEX-SSM, SSM, Chimera | FLEX-GreedyML, Hydra |
| Extended patches | FLEX-SSM, FLEX-MUSIC | Subspace-SBL+, FLEX-GreedyML |
| Low SNR | Subspace-SBL, FLEX-SSM | Subspace-SBL, Chimera |
| Unknown/general | FLEX-SSM, Chimera | FLEX-GreedyML |

1. **Focal Source**: A single point dipole at moderate-to-high SNR (3–5 dB). This baseline scenario tests pure localization accuracy and favors sparse/dipolar methods.
2. **Multi Source**: 2–4 simultaneously active focal dipoles at low-to-moderate SNR (1–3 dB). Tests the ability to resolve multiple concurrent sources without mutual interference.
3. **Extended Source**: 2–4 spatially extended contiguous Gaussian patches (spatial orders 1–3) at moderate SNR (1–3 dB). Uses the contiguous Gaussian patch model (Section 5.1) with rank-1 or rank-2 temporal dynamics within each patch, producing irregularly shaped cortical activations.
4. **Noisy**: 1–3 sources with variable spatial extent (orders 0–2) under challenging low-SNR conditions (−5 to 0 dB). Tests robustness to noise, which is the primary practical challenge in clinical and cognitive applications.

All four scenarios use the “realistic” noise model (Section 5.3) with a low-rank artifact dimension drawn uniformly from [2, 8].

### 6.3 Ranking Procedure

Solver rankings are computed from the frozen paper snapshot rather than re-derived manually in the manuscript. For each dataset, solvers receive dense ranks on four metrics: EMD, AP, MLE, and correlation. Spatial dispersion is reported but excluded from ranking. The per-dataset rank is the mean of those four metric ranks, and the global rank is the mean of the four per-dataset ranks. All ties and averages reported below follow that benchmark-runner definition.

### 6.4 Experimental Setup

The paper reports a frozen EEG benchmark snapshot containing 106 evaluated methods on a shared BioSemi-32 electrode montage with an `ico3` source space comprising *n* = 1,284 fixed-orientation dipoles. The snapshot covers 4 scenarios with 50 simulations per scenario. It therefore supports claims about this specific EEG benchmark family: synthetic EEG data, one 32-channel montage, one source space resolution, and one fixed-orientation forward-model configuration.

For each simulation, a noise covariance matrix is estimated from a separate 200-time-point baseline segment drawn independently from the same noise model (spatial covariance and temporal coloring) as the noise added to the signal. This simulates the common experimental practice of estimating noise statistics from an empty-room recording or pre-stimulus baseline. The empirical covariance is regularized with a fixed shrinkage toward a scaled identity (*γ* = 0.05). Each solver receives this estimated noise covariance and uses it to spatially whiten both the leadfield and the data via eigendecomposition-based whitening: the noise covariance is decomposed as **C**_*n*_ = **VΛV**^⊤^, and the whitener **W** = **Λ**^−1/2^**V**^⊤^ is applied, with rank truncation to avoid amplifying near-zero-variance directions. This ensures that all solvers operate on data with approximately identity noise covariance, placing them on equal footing regardless of the original noise structure.

The `BenchmarkRunner` class automates the evaluation pipeline: it generates simulated data for each scenario, applies each solver with its recommended depth-weighting parameters, computes all metrics, and aggregates results. The novel solvers were not tuned on the exact simulation instances used in the frozen evaluation; new random draws were generated for benchmarking. However, development and evaluation share the same broad simulation assumptions, so external validity remains limited to nearby EEG regimes rather than being claimable as a universal solver ranking.

### 6.5 Benchmark Results

Figure 4 summarizes the trade-off between reconstruction quality (mean EMD) and inference time across all 106 methods. The full leaderboard and per-scenario tables are moved to the Appendix (Tables 5 to 9).

**Figure 3:**
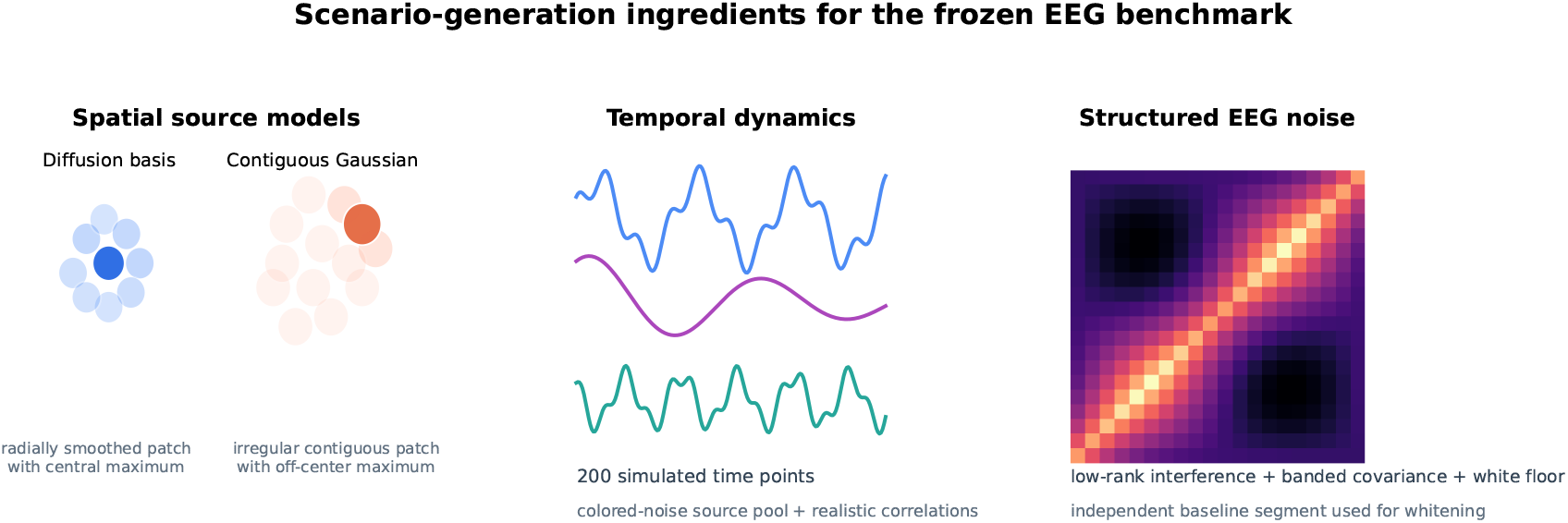
Scenario-generation overview for the frozen EEG benchmark. The benchmark combines two spatial source models, colored temporal dynamics, and structured EEG noise. The left panel illustrates the diffusion-basis and contiguous-Gaussian source constructions used in the simulation framework, without assigning them one-to-one to every scenario shown elsewhere in the manuscript.

**Figure 4:**
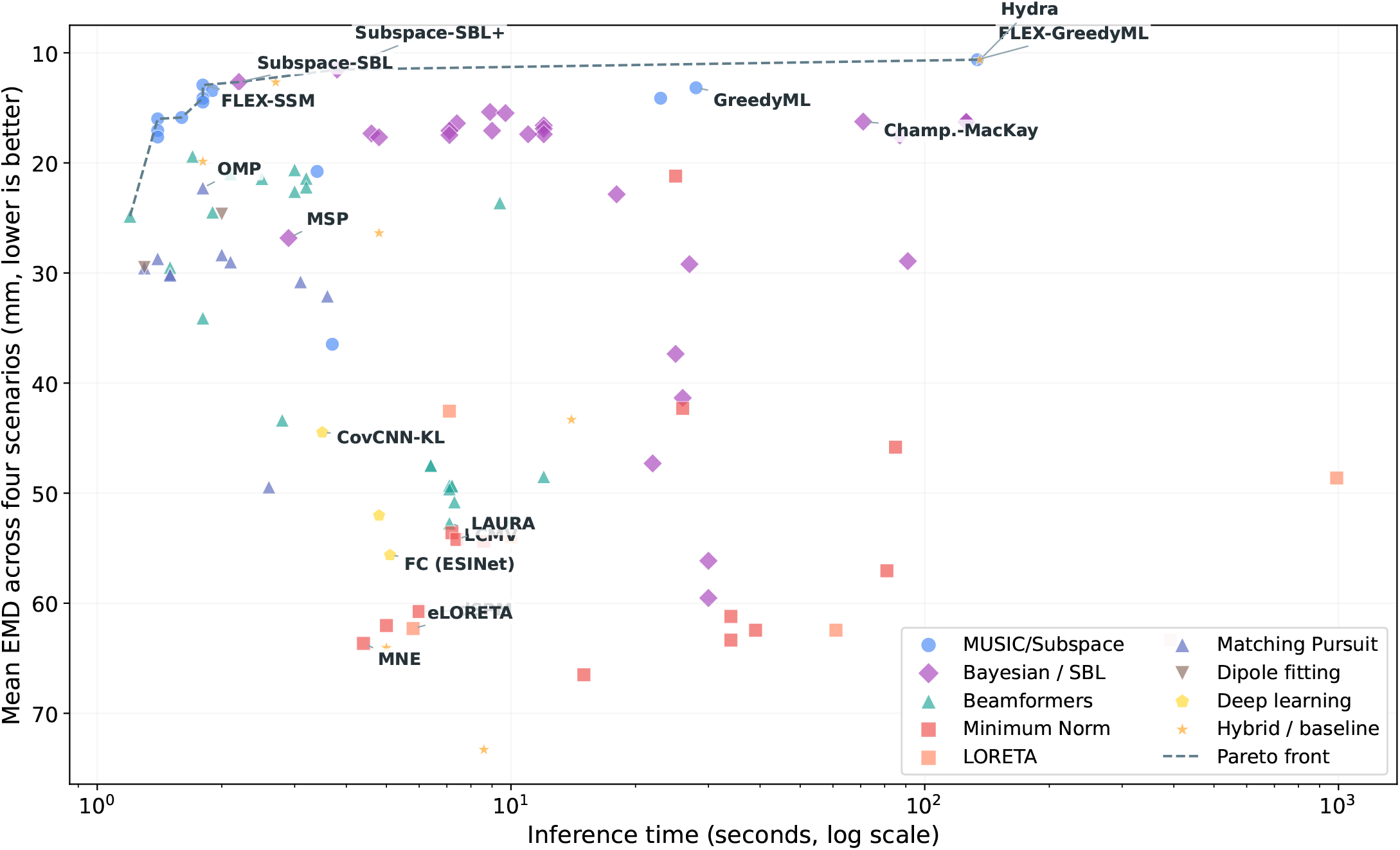
Performance–compute trade-off for the frozen EEG benchmark. The x-axis reports inference time only; neural-network training cost is discussed separately in the text. Labeled points include the Pareto front, canonical baselines, and the benchmarked neural-network baselines.

**Table 5:**
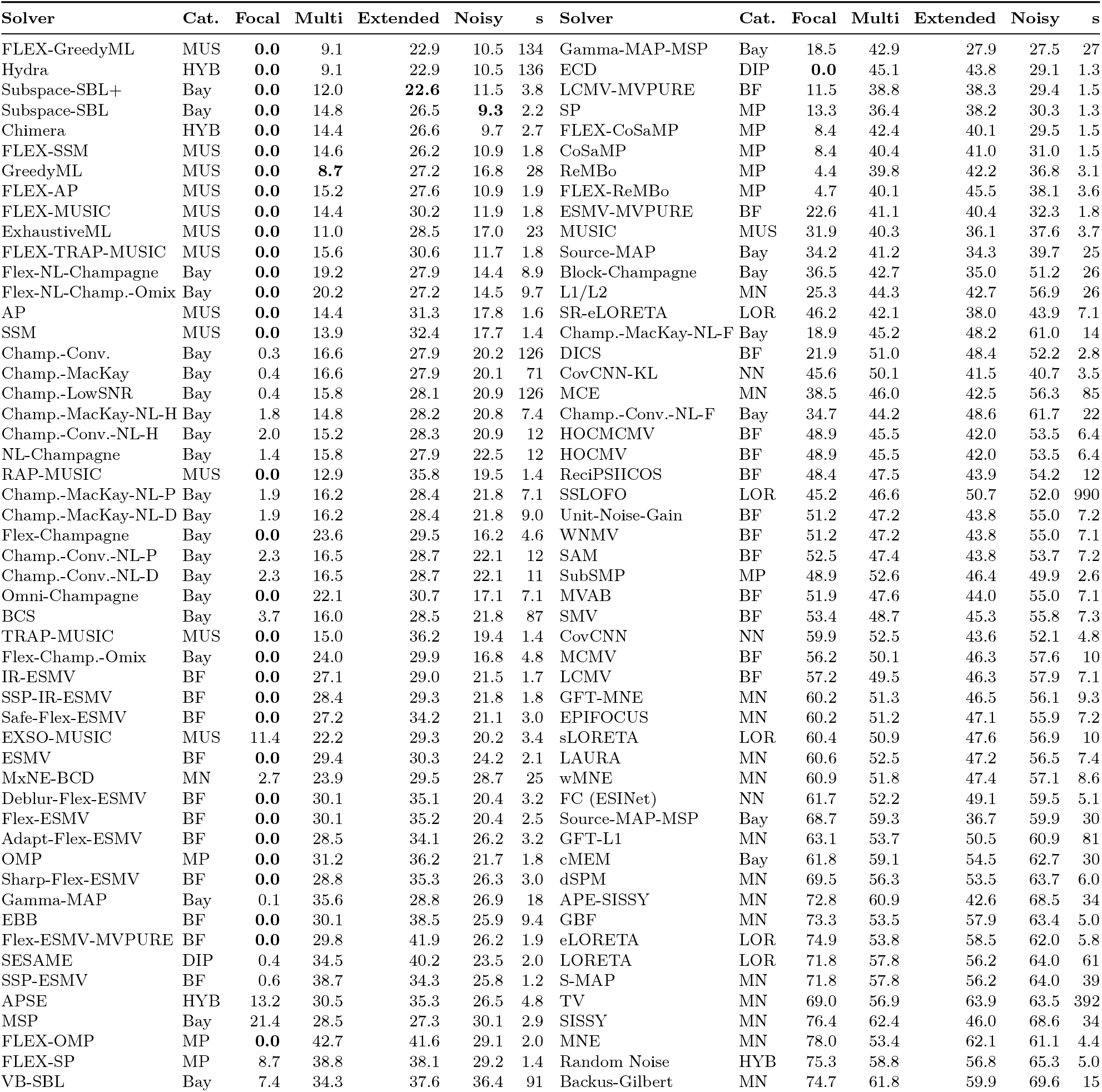
Earth mover’s distance (EMD in mm↓, lower is better) across all four EEG benchmark scenarios for all 106 evaluated solvers, sorted by mean EMD. Means are computed across 50 simulations per scenario. Average inference time in seconds excludes ANN training. Best value per scenario in **bold**.

**Table 6:** Results for the **Focal** scenario (means across 50 simulations, sorted by EMD). Best value per metric in **bold**.

| Solver | EMD (mm)↓ | AP↑ | MLE (mm)↓ | Corr↑ | Solver | EMD (mm)↓ | AP↑ | MLE (mm)↓ | Corr↑ |
| --- | --- | --- | --- | --- | --- | --- | --- | --- | --- |
| FLEX-GreedyML | **0.0** | **1.00** | **0.0** | **1.00** | CoSaMP | 8.4 | 0.88 | 6.4 | 0.73 |
| Hydra | **0.0** | **1.00** | **0.0** | **1.00** | FLEX-SP | 8.7 | 0.95 | 8.3 | 0.80 |
| Subspace-SBL+ | **0.0** | **1.00** | **0.0** | **1.00** | EXSO-MUSIC | 11.4 | 0.81 | 5.2 | 0.60 |
| Subspace-SBL | **0.0** | **1.00** | **0.0** | **1.00** | LCMV-MVPURE | 11.5 | 0.65 | 11.4 | 0.58 |
| Chimera | **0.0** | **1.00** | **0.0** | 0.98 | APSE | 13.2 | **1.00** | 6.1 | 0.53 |
| FLEX-SSM | **0.0** | **1.00** | **0.0** | **1.00** | SP | 13.3 | 0.94 | 11.1 | 0.77 |
| GreedyML | **0.0** | **1.00** | **0.0** | **1.00** | Gamma-MAP-MSP | 18.5 | 0.29 | 15.6 | 0.19 |
| FLEX-AP | **0.0** | **1.00** | **0.0** | **1.00** | Champ.-MacKay-NL-F | 18.9 | 0.80 | 19.0 | 0.72 |
| FLEX-MUSIC | **0.0** | **1.00** | **0.0** | **1.00** | MSP | 21.4 | 0.16 | 20.2 | 0.12 |
| ExhaustiveML | **0.0** | **1.00** | **0.0** | **1.00** | DICS | 21.9 | 0.80 | 16.6 | -0.01 |
| FLEX-TRAP-MUSIC | **0.0** | **1.00** | **0.0** | **1.00** | ESMV-MVPURE | 22.6 | 0.29 | 14.8 | -0.03 |
| Flex-NL-Champagne | **0.0** | **1.00** | **0.0** | **1.00** | L1/L2 | 25.3 | 0.51 | 25.3 | 0.45 |
| Flex-NL-Champ.-Omix | **0.0** | **1.00** | **0.0** | **1.00** | MUSIC | 31.9 | 0.18 | 22.0 | 0.21 |
| AP | **0.0** | **1.00** | **0.0** | **1.00** | Source-MAP | 34.2 | 0.51 | 15.4 | 0.24 |
| SSM | **0.0** | **1.00** | **0.0** | **1.00** | Champ.-Conv.-NL-F | 34.7 | 0.76 | 31.5 | 0.61 |
| RAP-MUSIC | **0.0** | **1.00** | **0.0** | **1.00** | Block-Champagne | 36.5 | 0.13 | 30.1 | 0.12 |
| Flex-Champagne | **0.0** | **1.00** | **0.0** | **1.00** | MCE | 38.5 | 0.29 | 34.7 | 0.25 |
| Omni-Champagne | **0.0** | **1.00** | **0.0** | **1.00** | SSLOFO | 45.2 | 0.41 | 18.5 | 0.14 |
| TRAP-MUSIC | **0.0** | **1.00** | **0.0** | **1.00** | CovCNN-KL | 45.6 | 0.41 | 27.7 | 0.28 |
| Flex-Champ.-Omix | **0.0** | **1.00** | **0.0** | **1.00** | SR-eLORETA | 46.2 | 0.64 | 16.2 | 0.14 |
| IR-ESMV | **0.0** | **1.00** | **0.0** | **1.00** | ReciPSIICOS | 48.4 | 0.72 | 13.3 | 0.18 |
| SSP-IR-ESMV | **0.0** | **1.00** | **0.0** | 0.99 | HOCMCMV | 48.9 | 0.23 | 22.7 | 0.09 |
| Safe-Flex-ESMV | **0.0** | **1.00** | 0.4 | 0.94 | HOCMV | 48.9 | 0.23 | 22.7 | 0.09 |
| ESMV | **0.0** | **1.00** | **0.0** | **1.00** | SubSMP | 48.9 | 0.51 | 24.9 | 0.49 |
| Deblur-Flex-ESMV | **0.0** | **1.00** | **0.0** | 0.98 | Unit-Noise-Gain | 51.2 | 0.38 | 15.8 | 0.09 |
| Flex-ESMV | **0.0** | **1.00** | **0.0** | 0.98 | WNMV | 51.2 | 0.38 | 15.8 | 0.09 |
| Adapt-Flex-ESMV | **0.0** | **1.00** | 0.5 | 0.94 | MVAB | 51.9 | 0.45 | 15.0 | 0.09 |
| OMP | **0.0** | **1.00** | **0.0** | **1.00** | SAM | 52.5 | 0.25 | 22.9 | 0.09 |
| Sharp-Flex-ESMV | **0.0** | **1.00** | 0.4 | 0.94 | SMV | 53.4 | 0.85 | 10.9 | 0.09 |
| EBB | **0.0** | **1.00** | **0.0** | **1.00** | MCMV | 56.2 | 0.06 | 25.6 | 0.05 |
| Flex-ESMV-MVPURE | **0.0** | **1.00** | **0.0** | **1.00** | LCMV | 57.2 | 0.18 | 22.8 | 0.08 |
| FLEX-OMP | **0.0** | **1.00** | **0.0** | **1.00** | MEM | 58.6 | 0.12 | 48.2 | 0.04 |
| ISubSMP | **0.0** | **1.00** | **0.0** | **1.00** | CovCNN | 59.9 | 0.21 | 39.1 | 0.10 |
| ECD | **0.0** | **1.00** | **0.0** | **1.00** | GFT-MNE | 60.2 | 0.05 | 35.0 | 0.06 |
| Gamma-MAP | 0.1 | **1.00** | 0.6 | 0.99 | EPIFOCUS | 60.2 | 0.11 | 38.9 | 0.08 |
| Champ.-Conv. | 0.3 | **1.00** | 0.7 | **1.00** | sLORETA | 60.4 | 0.43 | 21.5 | 0.08 |
| Champ.-MacKay | 0.4 | **1.00** | 0.9 | **1.00** | LAURA | 60.6 | 0.09 | 41.9 | 0.07 |
| Champ.-LowSNR | 0.4 | **1.00** | 1.6 | 0.99 | wMNE | 60.9 | 0.15 | 34.1 | 0.08 |
| SESAME | 0.4 | 0.97 | 0.4 | 0.97 | FC (ESINet) | 61.7 | 0.16 | 32.3 | 0.07 |
| SSP-ESMV | 0.6 | 0.97 | 0.5 | 0.97 | cMEM | 61.8 | 0.05 | 42.4 | 0.04 |
| NL-Champagne | 1.4 | **1.00** | 4.4 | 0.99 | GFT-L1 | 63.1 | 0.02 | 44.1 | 0.03 |
| Champ.-MacKay-NL-H | 1.8 | 0.95 | 4.1 | 0.92 | Source-MAP-MSP | 68.7 | 0.07 | 50.3 | 0.06 |
| Champ.-MacKay-NL-P | 1.9 | 0.92 | 5.4 | 0.91 | TV | 69.0 | 0.04 | 47.8 | 0.03 |
| Champ.-MacKay-NL-D | 1.9 | 0.92 | 5.4 | 0.91 | dSPM | 69.5 | 0.03 | 42.6 | 0.04 |
| Champ.-Conv.-NL-H | 2.0 | 0.93 | 4.9 | 0.92 | LORETA | 71.8 | 0.03 | 50.9 | 0.02 |
| Champ.-Conv.-NL-P | 2.3 | 0.92 | 5.6 | 0.90 | S-MAP | 71.8 | 0.03 | 50.9 | 0.02 |
| Champ.-Conv.-NL-D | 2.3 | 0.92 | 5.6 | 0.90 | APE-SISSY | 72.8 | 0.03 | 57.6 | 0.03 |
| MxNE-BCD | 2.7 | 0.94 | 4.4 | 0.92 | GBF | 73.3 | 0.03 | 56.9 | 0.04 |
| BCS | 3.7 | 0.93 | 7.6 | 0.91 | Backus-Gilbert | 74.7 | 0.01 | 64.9 | -0.00 |
| ReMBo | 4.4 | 0.99 | 12.5 | 0.97 | eLORETA | 74.9 | 0.03 | 64.0 | 0.04 |
| FLEX-ReMBo | 4.7 | 0.99 | 12.0 | 0.97 | Random Noise | 75.3 | 0.00 | 59.2 | -0.00 |
| VB-SBL | 7.4 | 0.80 | 6.1 | 0.75 | SISSY | 76.4 | 0.01 | 63.6 | 0.03 |
| FLEX-CoSaMP | 8.4 | 0.88 | 6.4 | 0.73 | MNE | 78.0 | 0.03 | 63.6 | 0.04 |

**Table 7:**
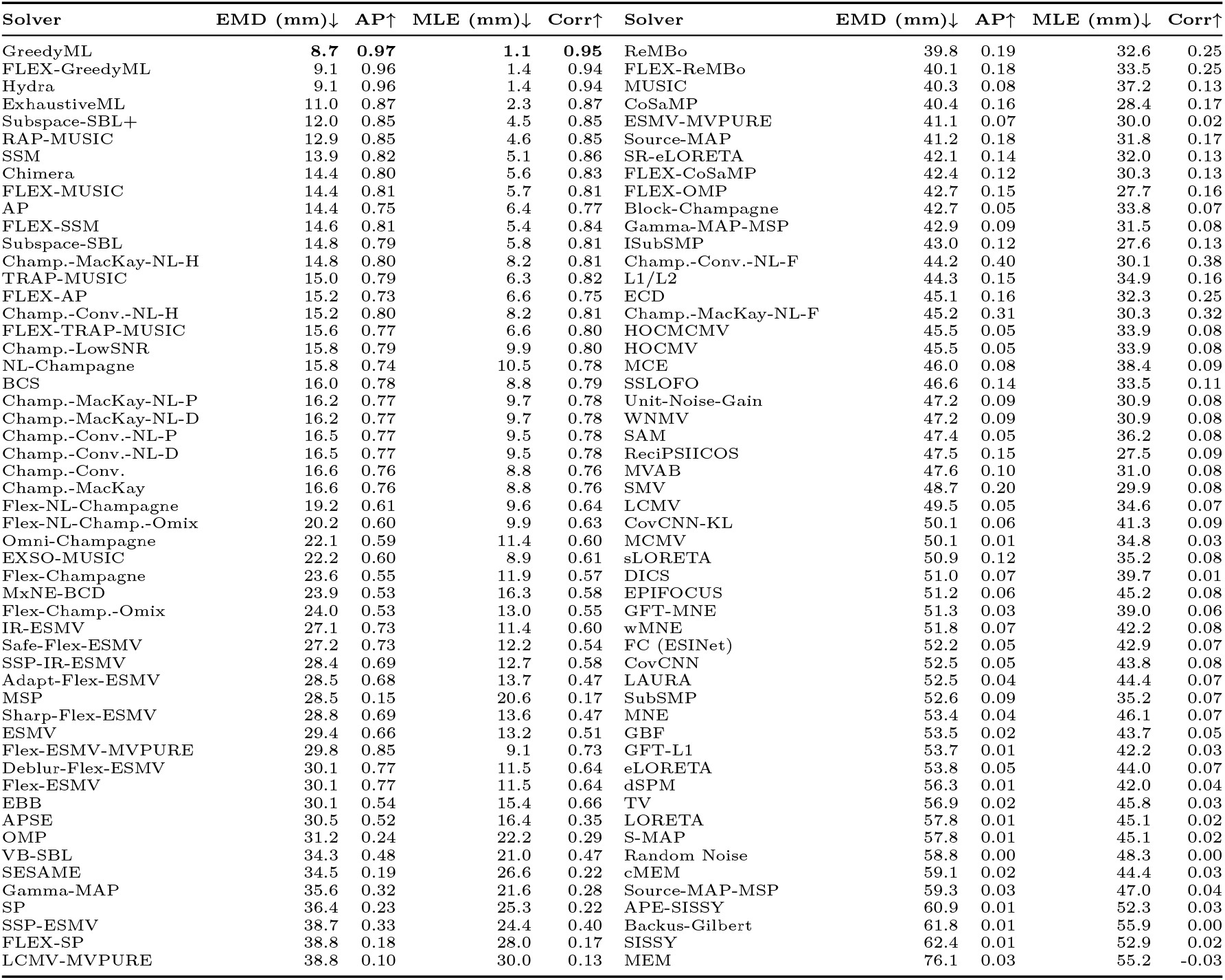
Results for the **Multi** scenario (means across 50 simulations, sorted by EMD). Best value per metric in **bold**.

**Table 8:** Results for the **Extended** scenario (means across 50 simulations, sorted by EMD). Best value per metric in **bold**.

| Solver | EMD (mm)↓ | AP↑ | MLE (mm)↓ | Corr↑ | Solver | EMD (mm)↓ | AP↑ | MLE (mm)↓ | Corr↑ |
| --- | --- | --- | --- | --- | --- | --- | --- | --- | --- |
| Subspace-SBL+ | **22.6** | 0.43 | 16.9 | 0.46 | FLEX-SP | 38.1 | 0.12 | 32.0 | 0.02 |
| FLEX-GreedyML | 22.9 | **0.45** | **13.8** | **0.53** | SP | 38.2 | 0.05 | 29.7 | 0.05 |
| Hydra | 22.9 | **0.45** | **13.8** | **0.53** | LCMV-MVPURE | 38.3 | 0.05 | 32.3 | 0.05 |
| FLEX-SSM | 26.2 | 0.40 | 17.8 | 0.47 | EBB | 38.5 | 0.08 | 27.6 | 0.16 |
| Subspace-SBL | 26.5 | 0.41 | 18.1 | 0.46 | FLEX-CoSaMP | 40.1 | 0.06 | 32.9 | 0.05 |
| Chimera | 26.6 | 0.34 | 19.4 | 0.41 | SESAME | 40.2 | 0.04 | 31.6 | 0.04 |
| GreedyML | 27.2 | 0.08 | 18.8 | 0.15 | ESMV-MVPURE | 40.4 | 0.04 | 32.8 | 0.01 |
| Flex-NL-Champ.-Omix | 27.2 | 0.38 | 20.9 | 0.41 | CoSaMP | 41.0 | 0.04 | 32.8 | 0.03 |
| MSP | 27.3 | 0.31 | 20.8 | 0.30 | ISubSMP | 41.1 | 0.10 | 31.0 | 0.04 |
| FLEX-AP | 27.6 | 0.36 | 19.6 | 0.42 | CovCNN-KL | 41.5 | 0.11 | 38.2 | 0.15 |
| Flex-NL-Champagne | 27.9 | 0.38 | 20.8 | 0.41 | FLEX-OMP | 41.6 | 0.11 | 30.5 | 0.04 |
| Champ.-Conv. | 27.9 | 0.10 | 20.8 | 0.17 | Flex-ESMV-MVPURE | 41.9 | 0.07 | 27.9 | 0.11 |
| Champ.-MacKay | 27.9 | 0.09 | 21.0 | 0.17 | HOCMCMV | 42.0 | 0.08 | 34.3 | 0.13 |
| NL-Champagne | 27.9 | 0.12 | 20.8 | 0.19 | HOCMV | 42.0 | 0.08 | 34.3 | 0.13 |
| Gamma-MAP-MSP | 27.9 | 0.40 | 18.9 | 0.32 | ReMBo | 42.2 | 0.04 | 36.1 | 0.06 |
| Champ.-LowSNR | 28.1 | 0.10 | 21.0 | 0.17 | MCE | 42.5 | 0.05 | 38.2 | 0.08 |
| Champ.-MacKay-NL-H | 28.2 | 0.10 | 21.5 | 0.17 | APE-SISSY | 42.6 | 0.11 | 39.5 | 0.18 |
| Champ.-Conv.-NL-H | 28.3 | 0.10 | 21.4 | 0.17 | L1/L2 | 42.7 | 0.06 | 35.7 | 0.09 |
| Champ.-MacKay-NL-P | 28.4 | 0.10 | 22.1 | 0.17 | CovCNN | 43.6 | 0.09 | 40.8 | 0.13 |
| Champ.-MacKay-NL-D | 28.4 | 0.10 | 22.1 | 0.17 | ECD | 43.8 | 0.04 | 35.9 | 0.06 |
| ExhaustiveML | 28.5 | 0.07 | 19.4 | 0.13 | Unit-Noise-Gain | 43.8 | 0.09 | 33.2 | 0.14 |
| BCS | 28.5 | 0.11 | 21.9 | 0.17 | WNMV | 43.8 | 0.09 | 33.2 | 0.14 |
| Champ.-Conv.-NL-P | 28.7 | 0.10 | 22.4 | 0.17 | SAM | 43.8 | 0.07 | 36.7 | 0.13 |
| Champ.-Conv.-NL-D | 28.7 | 0.10 | 22.4 | 0.17 | ReciPSIICOS | 43.9 | 0.09 | 30.9 | 0.10 |
| Gamma-MAP | 28.8 | 0.14 | 21.4 | 0.16 | MVAB | 44.0 | 0.08 | 33.1 | 0.13 |
| IR-ESMV | 29.0 | 0.15 | 25.6 | 0.21 | SMV | 45.3 | 0.11 | 32.5 | 0.14 |
| SSP-IR-ESMV | 29.3 | 0.14 | 26.7 | 0.20 | FLEX-ReMBo | 45.5 | 0.04 | 38.7 | 0.04 |
| EXSO-MUSIC | 29.3 | 0.26 | 23.4 | 0.33 | SISSY | 46.0 | 0.09 | 41.2 | 0.15 |
| Flex-Champagne | 29.5 | 0.37 | 23.6 | 0.37 | MCMV | 46.3 | 0.05 | 34.4 | 0.08 |
| MxNE-BCD | 29.5 | 0.09 | 23.7 | 0.16 | LCMV | 46.3 | 0.07 | 35.0 | 0.12 |
| Flex-Champ.-Omix | 29.9 | 0.36 | 23.3 | 0.37 | SubSMP | 46.4 | 0.10 | 30.0 | 0.02 |
| FLEX-MUSIC | 30.2 | 0.35 | 20.5 | 0.41 | GFT-MNE | 46.5 | 0.11 | 38.5 | 0.17 |
| ESMV | 30.3 | 0.14 | 26.4 | 0.18 | EPIFOCUS | 47.1 | 0.08 | 45.5 | 0.14 |
| FLEX-TRAP-MUSIC | 30.6 | 0.35 | 21.2 | 0.40 | LAURA | 47.2 | 0.08 | 42.3 | 0.14 |
| Omni-Champagne | 30.7 | 0.24 | 23.5 | 0.29 | wMNE | 47.4 | 0.09 | 42.3 | 0.14 |
| AP | 31.3 | 0.06 | 23.6 | 0.10 | sLORETA | 47.6 | 0.10 | 36.7 | 0.14 |
| SSM | 32.4 | 0.06 | 23.5 | 0.12 | Champ.-MacKay-NL-F | 48.2 | 0.05 | 36.9 | 0.07 |
| Adapt-Flex-ESMV | 34.1 | 0.22 | 25.1 | 0.25 | DICS | 48.4 | 0.04 | 38.8 | -0.00 |
| Safe-Flex-ESMV | 34.2 | 0.22 | 23.9 | 0.25 | Champ.-Conv.-NL-F | 48.6 | 0.06 | 37.7 | 0.08 |
| SSP-ESMV | 34.3 | 0.07 | 29.4 | 0.11 | FC (ESINet) | 49.1 | 0.11 | 40.3 | 0.18 |
| Source-MAP | 34.3 | 0.13 | 33.2 | 0.20 | GFT-L1 | 50.5 | 0.09 | 40.8 | 0.14 |
| Block-Champagne | 35.0 | 0.25 | 29.5 | 0.34 | SSLOFO | 50.7 | 0.08 | 44.5 | 0.12 |
| Deblur-Flex-ESMV | 35.1 | 0.19 | 23.8 | 0.23 | dSPM | 53.5 | 0.04 | 41.5 | 0.07 |
| Flex-ESMV | 35.2 | 0.22 | 23.8 | 0.25 | cMEM | 54.5 | 0.06 | 43.1 | 0.08 |
| Sharp-Flex-ESMV | 35.3 | 0.20 | 25.1 | 0.22 | LORETA | 56.2 | 0.07 | 44.1 | 0.09 |
| APSE | 35.3 | 0.15 | 26.5 | 0.11 | S-MAP | 56.2 | 0.07 | 44.1 | 0.09 |
| RAP-MUSIC | 35.8 | 0.06 | 26.7 | 0.10 | Random Noise | 56.8 | 0.03 | 48.3 | 0.00 |
| MUSIC | 36.1 | 0.08 | 38.0 | 0.14 | GBF | 57.9 | 0.05 | 51.3 | 0.10 |
| TRAP-MUSIC | 36.2 | 0.06 | 27.2 | 0.10 | eLORETA | 58.5 | 0.04 | 56.6 | 0.06 |
| OMP | 36.2 | 0.04 | 28.1 | 0.07 | Backus-Gilbert | 59.9 | 0.04 | 54.8 | 0.01 |
| Source-MAP-MSP | 36.7 | 0.11 | 34.0 | 0.15 | MNE | 62.1 | 0.04 | 55.6 | 0.06 |
| VB-SBL | 37.6 | 0.15 | 27.4 | 0.20 | TV | 63.9 | 0.04 | 54.4 | 0.08 |
| SR-eLORETA | 38.0 | 0.10 | 33.4 | 0.16 | MEM | 72.6 | 0.06 | 54.2 | -0.05 |

**Table 9:** Results for the **Noisy** scenario (means across 50 simulations, sorted by EMD). Best value per metric in **bold**.

| Solver | EMD (mm)↓ | AP↑ | MLE (mm)↓ | Corr↑ | Solver | EMD (mm)↓ | AP↑ | MLE (mm)↓ | Corr↑ |
| --- | --- | --- | --- | --- | --- | --- | --- | --- | --- |
| Subspace-SBL | **9.3** | **0.88** | **3.9** | **0.90** | FLEX-CoSaMP | 29.5 | 0.22 | 22.4 | 0.19 |
| Chimera | 9.7 | **0.88** | 4.2 | **0.90** | MSP | 30.1 | 0.27 | 24.7 | 0.23 |
| FLEX-GreedyML | 10.5 | **0.88** | 4.0 | **0.90** | SP | 30.3 | 0.21 | 19.9 | 0.24 |
| Hydra | 10.5 | **0.88** | 4.0 | **0.90** | CoSaMP | 31.0 | 0.19 | 22.9 | 0.19 |
| FLEX-SSM | 10.9 | **0.88** | 4.0 | **0.90** | ISubSMP | 31.9 | 0.43 | 21.1 | 0.20 |
| FLEX-AP | 10.9 | 0.86 | 4.3 | 0.87 | ESMV-MVPURE | 32.3 | 0.10 | 23.8 | 0.03 |
| Subspace-SBL+ | 11.5 | 0.87 | 5.7 | 0.87 | VB-SBL | 36.4 | 0.25 | 22.4 | 0.28 |
| FLEX-TRAP-MUSIC | 11.7 | 0.86 | 5.1 | 0.87 | ReMBo | 36.8 | 0.17 | 33.9 | 0.24 |
| FLEX-MUSIC | 11.9 | 0.87 | 4.8 | 0.87 | MUSIC | 37.6 | 0.07 | 33.6 | 0.14 |
| Flex-NL-Champagne | 14.4 | 0.81 | 8.0 | 0.80 | FLEX-ReMBo | 38.1 | 0.16 | 37.0 | 0.22 |
| Flex-NL-Champ.-Omix | 14.5 | 0.81 | 8.4 | 0.81 | Source-MAP | 39.7 | 0.13 | 30.7 | 0.17 |
| Flex-Champagne | 16.2 | 0.80 | 8.7 | 0.78 | CovCNN-KL | 40.7 | 0.10 | 31.4 | 0.13 |
| GreedyML | 16.8 | 0.36 | 9.1 | 0.54 | SR-eLORETA | 43.9 | 0.10 | 32.8 | 0.14 |
| Flex-Champ.-Omix | 16.8 | 0.79 | 9.0 | 0.78 | SubSMP | 49.9 | 0.27 | 32.8 | 0.06 |
| ExhaustiveML | 17.0 | 0.36 | 9.2 | 0.54 | Block-Champagne | 51.2 | 0.17 | 38.4 | 0.17 |
| Omni-Champagne | 17.1 | 0.66 | 10.1 | 0.68 | SSLOFO | 52.0 | 0.12 | 29.7 | 0.11 |
| SSM | 17.7 | 0.34 | 9.3 | 0.52 | CovCNN | 52.1 | 0.05 | 40.8 | 0.09 |
| AP | 17.8 | 0.34 | 9.6 | 0.52 | DICS | 52.2 | 0.09 | 38.0 | -0.01 |
| TRAP-MUSIC | 19.4 | 0.33 | 11.4 | 0.52 | HOCMCMV | 53.5 | 0.07 | 33.1 | 0.09 |
| RAP-MUSIC | 19.5 | 0.34 | 11.3 | 0.52 | HOCMV | 53.5 | 0.07 | 33.1 | 0.09 |
| Champ.-MacKay | 20.1 | 0.35 | 12.2 | 0.49 | SAM | 53.7 | 0.05 | 38.0 | 0.09 |
| Champ.-Conv. | 20.2 | 0.35 | 12.3 | 0.49 | ReciPSIICOS | 54.2 | 0.11 | 28.3 | 0.07 |
| EXSO-MUSIC | 20.2 | 0.41 | 14.3 | 0.41 | Unit-Noise-Gain | 55.0 | 0.10 | 28.4 | 0.09 |
| Deblur-Flex-ESMV | 20.4 | 0.51 | 9.9 | 0.68 | WNMV | 55.0 | 0.10 | 28.4 | 0.09 |
| Flex-ESMV | 20.4 | 0.67 | 10.0 | 0.69 | MVAB | 55.0 | 0.11 | 27.8 | 0.09 |
| Champ.-MacKay-NL-H | 20.8 | 0.34 | 16.8 | 0.47 | SMV | 55.8 | 0.16 | 27.2 | 0.09 |
| Champ.-LowSNR | 20.9 | 0.36 | 14.0 | 0.49 | EPIFOCUS | 55.9 | 0.05 | 46.7 | 0.09 |
| Champ.-Conv.-NL-H | 20.9 | 0.34 | 17.1 | 0.47 | GFT-MNE | 56.1 | 0.07 | 38.9 | 0.09 |
| Safe-Flex-ESMV | 21.1 | 0.62 | 15.6 | 0.56 | MCE | 56.3 | 0.02 | 45.9 | 0.04 |
| IR-ESMV | 21.5 | 0.40 | 14.3 | 0.46 | LAURA | 56.5 | 0.05 | 45.5 | 0.08 |
| OMP | 21.7 | 0.26 | 15.4 | 0.40 | L1/L2 | 56.9 | 0.06 | 42.8 | 0.10 |
| Champ.-MacKay-NL-P | 21.8 | 0.33 | 17.9 | 0.46 | sLORETA | 56.9 | 0.08 | 35.0 | 0.09 |
| Champ.-MacKay-NL-D | 21.8 | 0.33 | 17.9 | 0.46 | wMNE | 57.1 | 0.07 | 39.7 | 0.09 |
| BCS | 21.8 | 0.34 | 17.3 | 0.47 | MCMV | 57.6 | 0.02 | 33.3 | 0.04 |
| SSP-IR-ESMV | 21.8 | 0.39 | 15.2 | 0.45 | LCMV | 57.9 | 0.04 | 37.0 | 0.08 |
| Champ.-Conv.-NL-P | 22.1 | 0.33 | 18.8 | 0.46 | FC (ESINet) | 59.5 | 0.06 | 45.7 | 0.08 |
| Champ.-Conv.-NL-D | 22.1 | 0.33 | 18.8 | 0.46 | Source-MAP-MSP | 59.9 | 0.03 | 46.6 | 0.05 |
| NL-Champagne | 22.5 | 0.28 | 20.8 | 0.42 | GFT-L1 | 60.9 | 0.03 | 45.5 | 0.05 |
| SESAME | 23.5 | 0.24 | 17.0 | 0.36 | Champ.-MacKay-NL-F | 61.0 | 0.07 | 42.6 | 0.10 |
| ESMV | 24.2 | 0.37 | 16.4 | 0.37 | MNE | 61.1 | 0.04 | 48.4 | 0.07 |
| SSP-ESMV | 25.8 | 0.29 | 19.8 | 0.40 | Champ.-Conv.-NL-F | 61.7 | 0.10 | 42.7 | 0.13 |
| EBB | 25.9 | 0.28 | 16.6 | 0.42 | eLORETA | 62.0 | 0.03 | 49.6 | 0.06 |
| Adapt-Flex-ESMV | 26.2 | 0.58 | 17.6 | 0.44 | cMEM | 62.7 | 0.04 | 45.2 | 0.05 |
| Flex-ESMV-MVPURE | 26.2 | 0.34 | 14.7 | 0.45 | GBF | 63.4 | 0.04 | 50.4 | 0.07 |
| Sharp-Flex-ESMV | 26.3 | 0.58 | 17.7 | 0.44 | TV | 63.5 | 0.04 | 50.9 | 0.05 |
| APSE | 26.5 | 0.38 | 17.9 | 0.29 | dSPM | 63.7 | 0.01 | 48.0 | 0.04 |
| Gamma-MAP | 26.9 | 0.29 | 19.4 | 0.30 | LORETA | 64.0 | 0.03 | 48.0 | 0.03 |
| Gamma-MAP-MSP | 27.5 | 0.41 | 21.6 | 0.26 | S-MAP | 64.0 | 0.03 | 48.0 | 0.03 |
| MxNE-BCD | 28.7 | 0.29 | 23.0 | 0.40 | Random Noise | 65.3 | 0.01 | 52.5 | 0.00 |
| FLEX-OMP | 29.1 | 0.45 | 20.4 | 0.21 | APE-SISSY | 68.5 | 0.02 | 59.8 | 0.04 |
| ECD | 29.1 | 0.23 | 20.5 | 0.37 | SISSY | 68.6 | 0.01 | 59.4 | 0.03 |
| FLEX-SP | 29.2 | 0.38 | 21.4 | 0.13 | Backus-Gilbert | 69.6 | 0.01 | 64.4 | 0.00 |
| LCMV-MVPURE | 29.4 | 0.16 | 23.0 | 0.19 | MEM | 85.9 | 0.02 | 68.6 | -0.03 |

**Table 10:** Complete solver reference (Part 1). A machine-readable version with module paths and provenance is provided in supplement/solver_reference.csv.

| Category | Solver | Canonical id / aliases |
| --- | --- | --- |
| Minimum Norm | mne | mne; none |
|  | gft-mne | gft-mne; none |
|  | wmne | wmne; none |
|  | dspm | dspm; none |
|  | MCE | mce; l1, fista, minimum current estimate |
|  | l1l2 | l1l2; mce-l1l2 |
|  | MxNE-BCD | mxne-bcd; mxne, mixed-norm-bcd |
|  | gft-l1 | gft-l1; gft-minimum-l1, gftmce |
|  | laura | laura; laur |
|  | backus-gilbert | backus-gilbert; b-g, bg |
|  | s-map | s-map; smap |
|  | tv | tv; total-variation, graph-tv |
|  | sissy | sissy; none |
|  | apesissy | apesissy; ape-sissy |
|  | GBF | gbf; basisfunctions, basis-functions |
|  | epifocus | epifocus; none |
| LORETA | self-regularized-eloreta | self-regularized-eloreta; sr-eloreta, sreloreta, selfregeloreta |
|  | loreta | loreta; lor |
|  | sloreta | sloreta; slor |
|  | eloreta | eloreta; elor |
|  | sslofo | sslofo; none |
| Bayesian / SBL | MacKay-Champagne | champagne-mackay; champagne, mackaychampagne, mackay-champagne, mcc |
|  | Convexity-Champagne | champagne-convexity; convexitychampagne, convexity-champagne, coc, champagne-em, emchampagne, em-champagne, emc, mm-champagne |
|  | TEM-Champagne | champagne-tem; temchampagne, tem-champagne, temc, t-em-champagne |
|  | AR-EM-Champagne | champagne-ar-em; aremchampagne, arem-champagne, aremc, ar-em-champagne |
|  | Low-SNR-Champagne | champagne-low-snr; lowsnrchampagne, low-snr-champagne, lowsnr-champagne, lsc |
|  | MacKay-Champagne (homoscedastic noise) | champagne-mackay-nl-homoscedastic; none |
|  | MacKay-Champagne (diag noise) | champagne-mackay-nl-diag; none |
|  | MacKay-Champagne (full noise) | champagne-mackay-nl-full; none |
|  | MacKay-Champagne (prec noise) | champagne-mackay-nl-prec; none |
|  | Convexity-Champagne (homoscedastic noise) | champagne-convexity-nl-homoscedastic; champagne-em-nl-homoscedastic |
|  | Convexity-Champagne (diag noise) | champagne-convexity-nl-diag; champagne-em-nl-diag |
|  | Convexity-Champagne (full noise) | champagne-convexity-nl-full; champagne-em-nl-full |
|  | Convexity-Champagne (prec noise) | champagne-convexity-nl-prec; champagne-em-nl-prec |
|  | nl-champagne | nl-champagne; nlchampagne, nl-champagne, nlc |
|  | omni-champagne | omni-champagne; omnichampagne, omni-champagne, omni champagne, oc |
|  | flex-champagne | flex-champagne; flexchampagne, flex-champagne, fc-champ |
|  | Flex-Champagne (ongoing-mixture tuned) | flex-champagne-omix; fc-omix |
|  | flex-nl-champagne | flex-nl-champagne; flexnlchampagne, flex-nl-champagne, fnlc |
|  | Flex-NL-Champagne (ongoing-mixture tuned) | flex-nl-champagne-omix; fnlc-omix |
|  | Block-Champagne | block-champagne; blockchampagne, block-champagne, bc-champ |
|  | gamma-map | gamma-map; gmap |
|  | source-map | source-map; none |
|  | gamma-map-msp | gamma-map-msp; none |
|  | source-map-msp | source-map-msp; none |
|  | msp | msp; multiple-sparse-priors |
|  | mem | mem; none |
|  | cmem | cmem; none |
|  | subspace-sbl | subspace-sbl; ssm-nlc, subspacesbl |
|  | subspace-sbl-plus | subspace-sbl-plus; ssm-nlc-plus, subspacesblplus |
|  | vb-sbl | vb-sbl; vbsbl, variational-bayes-sbl |
|  | bcs | bcs; none |

**Table 11:**
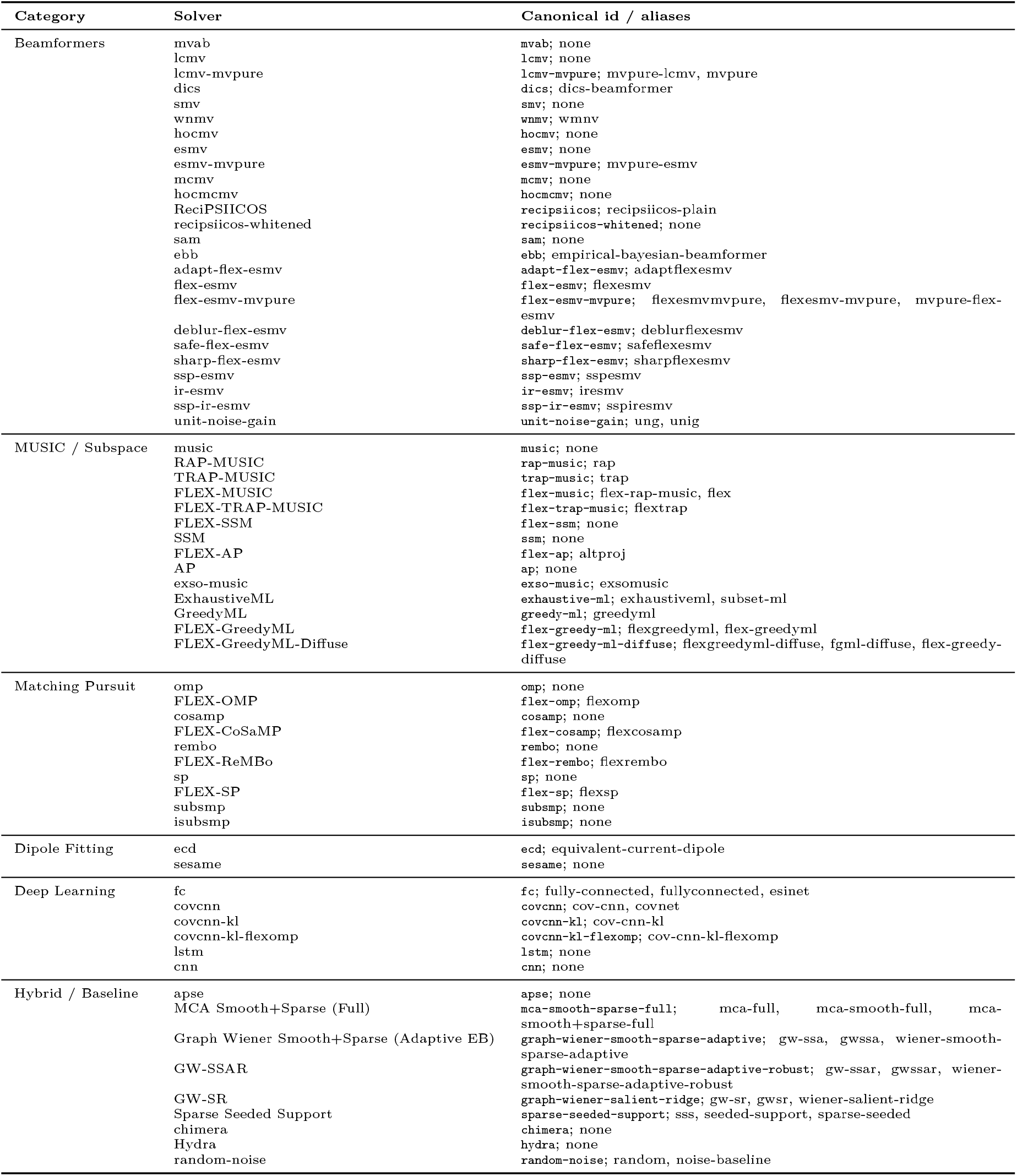
Complete solver reference (Part 2). A machine-readable version with module paths and provenance is provided in supplement/solver_reference.csv.

#### Overall ranking

The frozen snapshot yields a tie for best global rank between **FLEX-GreedyML** and **Hydra** (mean global rank 1.5), followed by **Subspace-SBL+** (4.0), **FLEX-SSM** (4.06), and **Subspace-SBL** (4.62). This top tier is dominated by flexible subspace scanners, hybrid switching methods, and Bayesian refinements built on subspace detection. The practical implication is not that a single universal winner exists, but that methods combining explicit source enumeration with flexible spatial extent modeling perform strongly across this benchmark family.

#### Focal and multi-source scenarios

The focal scenario is deliberately easy for accurate sparse scanners: many methods achieve essentially perfect localization, so speed and robustness become the main differentiators. The multi-source scenario is more informative. Here **GreedyML** yields the best mean EMD (8.7 mm), followed by **FLEX-GreedyML** and **Hydra** (9.1 mm), **ExhaustiveML** (11.0 mm), and **Subspace-SBL+** (12.0 mm). This is the regime in which explicit source-enumeration methods separate most clearly from diffuse minimum-norm solutions.

#### Extended source scenario

For spatially extended sources, **Subspace-SBL+** leads (EMD 22.6 mm), followed closely by **FLEX-GreedyML** and **Hydra** (22.9 mm), **FLEX-SSM** (26.2 mm), and **Subspace-SBL** (26.5 mm). This scenario uses the contiguous Gaussian patch model rather than the diffusion basis used in focal and multi-source conditions, which helps test whether flexible solvers generalize beyond a single source-construction rule. MSP also ranks well here, consistent with the advantage of distributed priors when the ground truth is genuinely extended. The qualitative comparison in Figure 1 illustrates the same point at the single-example level: for these moderately compact patches, subspace and Bayesian methods recover the bilateral generators more faithfully than the diffuse minimum-norm family. This figure should not be overgeneralized to all extended activity, however. If the true source prior were much broader or smoother than the contiguous Gaussian patches used here, methods such as MNE or eLORETA could become more competitive because their spatial smoothness assumptions would better match the generating process.

#### Noisy scenario

Under low-SNR conditions, **Subspace-SBL** achieves the best EMD (9.3 mm), closely followed by **Chimera** (9.7 mm), **FLEX-GreedyML** and **Hydra** (10.5 mm), and **FLEX-AP** and **FLEX-SSM** (10.9 mm). Bayesian methods generally maintain their performance better than beamformers under noise, consistent with the regularizing effect of their probabilistic priors.

#### Deep learning solvers

Three neural network solvers were included in the benchmark: CovCNN-KL, CovCNN, and FC. CovCNN-KL achieves the best performance among them (mean global rank 68.5, mean EMD 44.5 mm), placing it in the range of mid-tier classical beamformers such as DICS and LCMV. CovCNN (78.06, 52.0 mm) and FC (79.94, 55.6 mm) perform similarly to lower-ranked minimum norm methods. Notably, all three neural network solvers outperform the random-noise baseline and several classical methods, but fall substantially short of the top-performing classical solvers. The benchmark used reduced model capacity (24 dense units, 200 training epochs) to keep training tractable within the evaluation framework, resulting in approximately 10^6^ trainable parameters per model. These ANN results should therefore be interpreted as constrained-budget baselines rather than as a definitive upper bound on deep-learning performance.

## 7 Usage Examples

### 7.1 Basic Source Imaging

The following example demonstrates the core workflow:

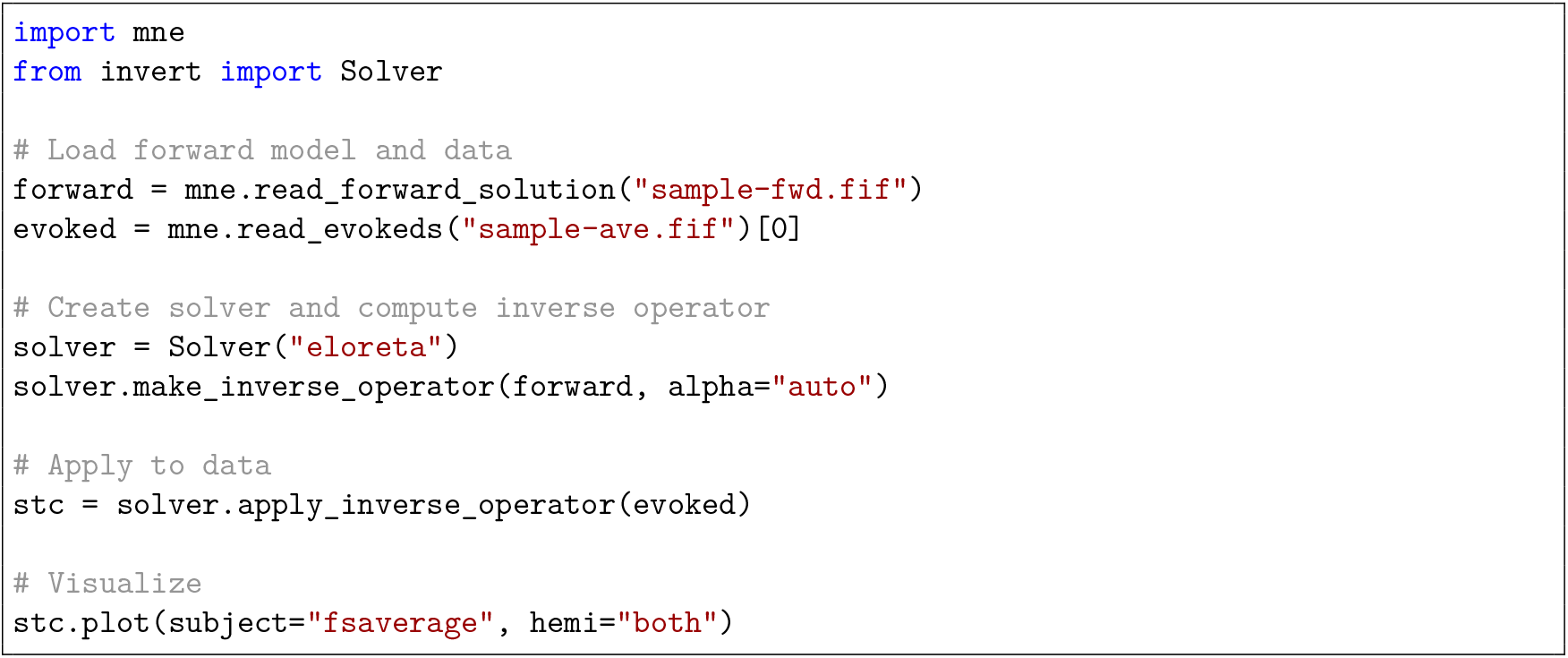

Switching to any other solver requires changing only the name string:

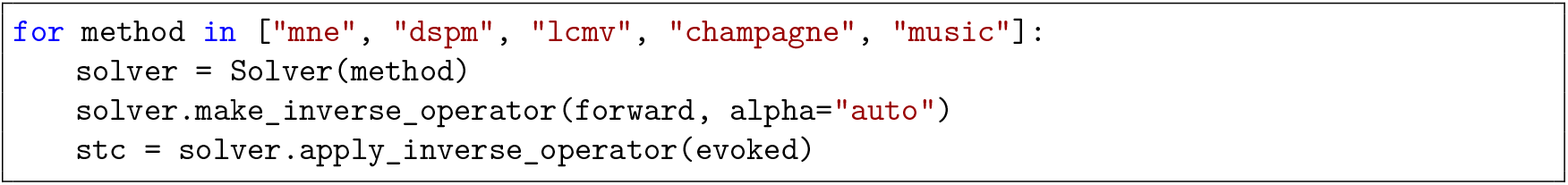

### 7.2 Simulation and Benchmarking

The same API exposes the simulation machinery used to build the benchmark:

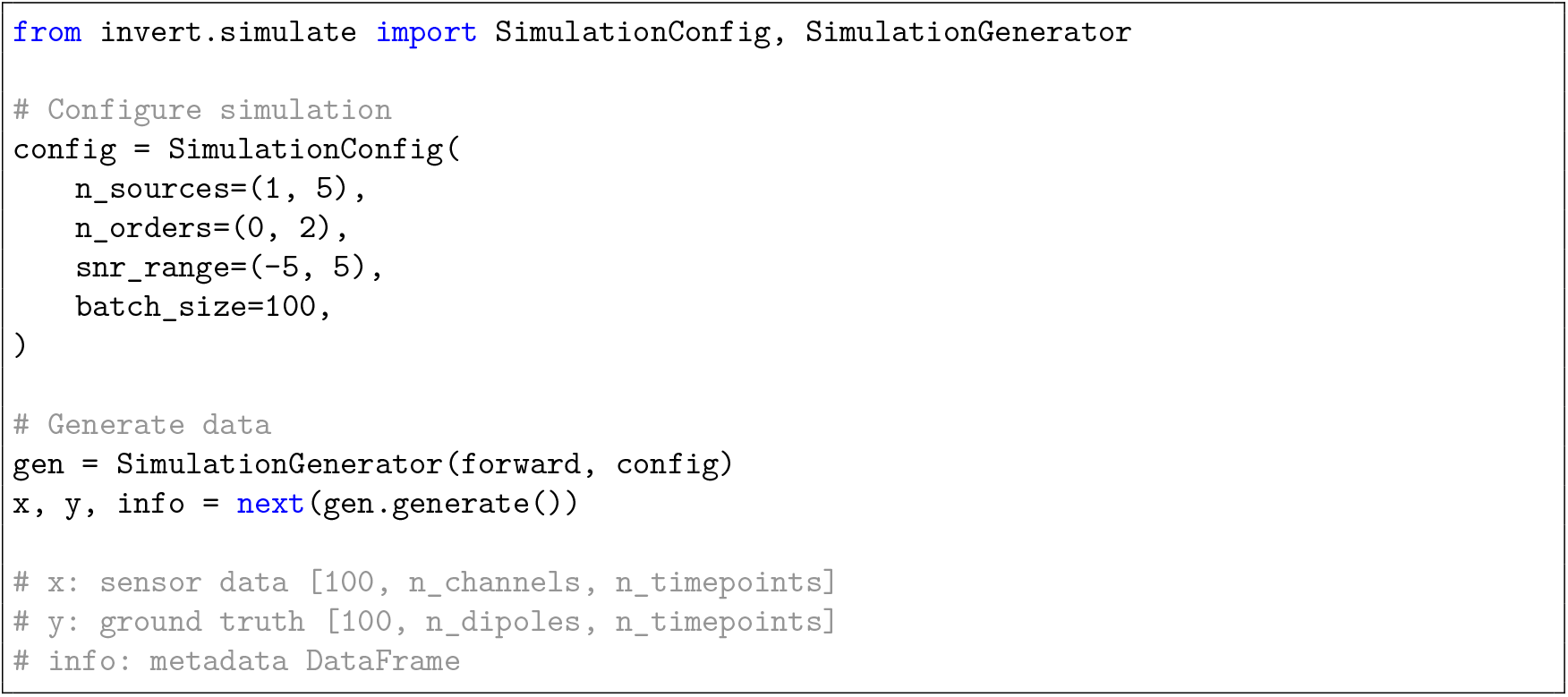

Figure 5 shows one representative sample from each benchmark scenario together with reconstructions from four solvers spanning different method families.

**Figure 5:**
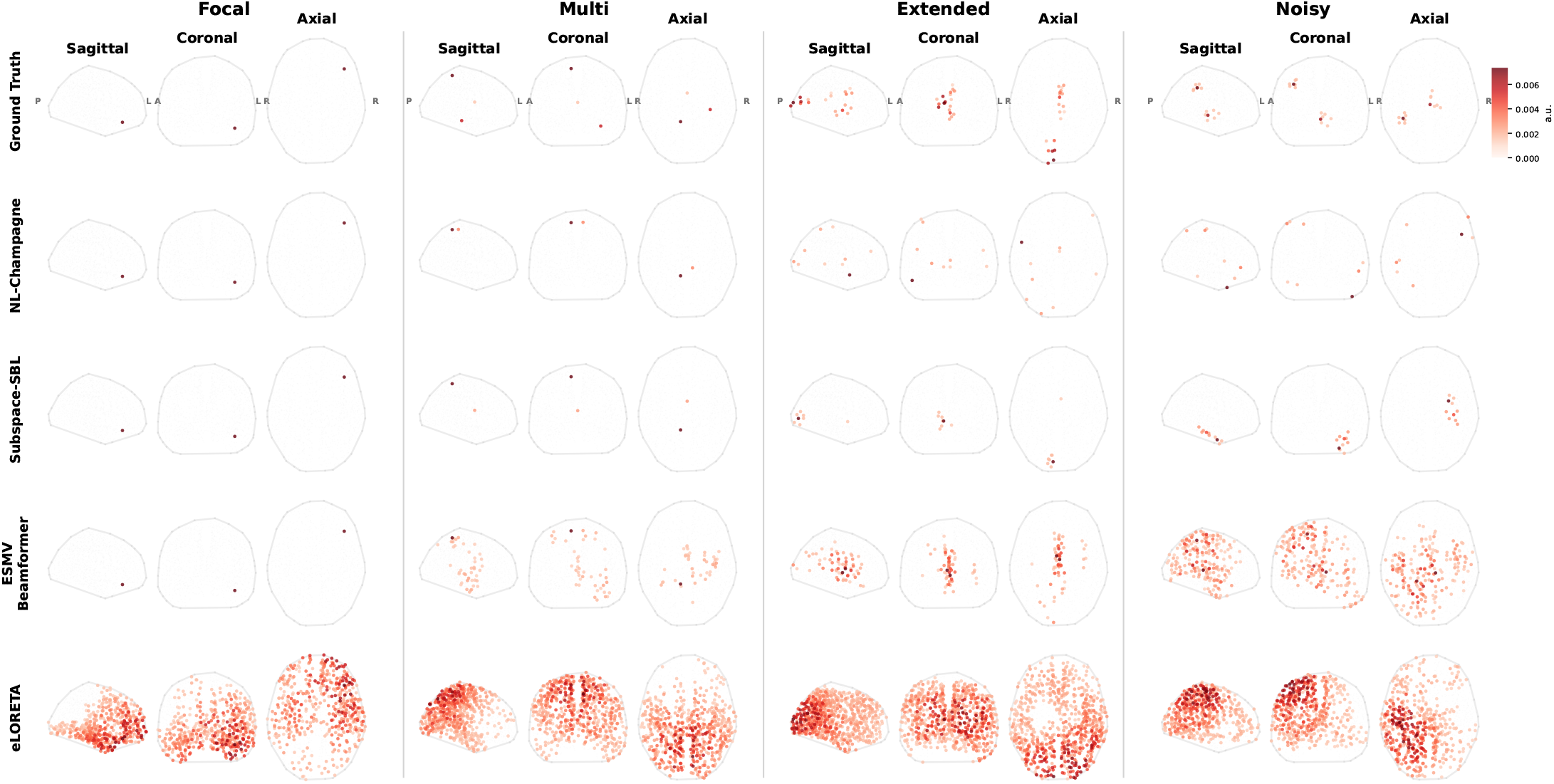
Qualitative reconstruction grid for one representative sample from each benchmark scenario. Columns show focal, multi-source, extended, and noisy simulations; rows show the ground truth together with reconstructions from NL-Champagne, Subspace-SBL, ESMV Beamformer, and eLORETA. The figure highlights distinct failure modes across scenarios: compact Bayesian solutions remain sparse in focal settings, ESMV is accurate for focal activity but becomes increasingly diffuse in the more challenging regimes, and eLORETA consistently produces broader spatial estimates.

## 8 Discussion

### 8.1 Practical Guide to Solver Selection

A key goal of this benchmark is to provide practical guidance for practitioners. Based on the results across all four scenarios, we offer the following recommendations, organized by the expected source configuration. Table 4 summarizes these recommendations.

#### When sources are expected to be focal

Many solvers achieve near-perfect performance on focal sources, so the differentiator is computational cost. Subspace methods (FLEX-MUSIC, FLEX-SSM, SSM) and matching pursuit methods (OMP) are fast (∼1−2 s) and highly accurate. Bayesian methods (Omni-Champagne, Flex-Champagne) achieve slightly better correlation but at higher computational cost (∼3−5 s).

#### When multiple sources are expected

Resolving multiple concurrent sources is the core strength of subspace scanning methods. FLEX-GreedyML and Hydra consistently lead, with SSM and ExhaustiveML close behind. Classical beamformers struggle in this regime due to signal cancellation between correlated sources.

#### When sources are spatially extended

Extended sources violate the sparsity assumption of many methods. Subspace-SBL+ and Subspace-SBL, which learn flexible source priors, excel here. Gamma-MAP-MSP, which employs a multiple sparse priors basis, also performs well (AP = 0.45). Pure sparse methods (OMP, CoSaMP, MCE) should be avoided.

#### When SNR is low

Subspace-SBL stands out under noisy conditions, consistently achieving the best rank across all metrics. Bayesian methods with noise learning (Champagne variants with homoscedastic or diagonal noise estimation) maintain reasonable performance. Beamformers and minimum norm methods degrade substantially as they lack the regularizing effect of structured source priors.

#### When no prior information is available

For a general-purpose solver that performs strongly across all scenarios in this benchmark, **FLEX-GreedyML** offers the best overall accuracy but at a higher computational cost (∼134 s per sample). For a fast default, **FLEX-SSM** (∼1.8 s) or **Chimera** (∼2.7 s) provide a good balance of performance and speed. Under this benchmark configuration, classical minimum norm defaults such as MNE and eLORETA rank well below several subspace, hybrid, and Bayesian alternatives.

#### Inference time

Figure 4 shows the performance–compute trade-off across all solvers (inference time only; training time for neural networks is excluded). Solver inference times span three orders of magnitude. The fastest solvers (< 2 s) include most subspace scanners (SSM, FLEX-SSM, AP, RAP-MUSIC), matching pursuit methods (OMP, CoSaMP), and simple beamformers (SSP-ESMV, LCMV-MVPURE). Moderate-cost solvers (2–10 s) include most Champagne variants, the ESMV family, and basic minimum norm methods. The most expensive solvers (> 30 s) include iterative Bayesian methods (VB-SBL: 91 s, BCS: 87 s), *L*_1_-based methods (MCE: 85 s), and hybrid methods (Hydra: 136 s, FLEX-GreedyML: 134 s). Total Variation (392 s) and SSLOFO (991 s) are the slowest solvers. Deep learning solvers have modest inference times (FC: ∼5 s, CovCNN-KL: ∼4 s) but require substantial upfront training (∼250−1,900 s per scenario). The Pareto front (Figure 4) is dominated by subspace methods at the fast end and FLEX-GreedyML at the accurate end, with Subspace-SBL variants offering an attractive middle ground.

### 8.2 Related Software

Several established toolboxes provide M/EEG source imaging functionality. `MNE-Python` [14] includes MNE, dSPM, sLORETA, eLORETA, LCMV, DICS, MUSIC, and RAP-MUSIC. Brainstorm [15] (MATLAB) provides minimum norm variants, beamformers, and dipole fitting. FieldTrip [16] (MATLAB) focuses on beamforming methods. SPM/DAiSS [17] (MATLAB) offers Bayesian approaches including MSP and variational methods. ESINet [12] introduced deep learning for EEG source imaging but was limited to fully connected architectures.

`invertmeeg` differs from these tools in scope and methodology. With 118 registered implementations and 106 methods included in the frozen EEG snapshot, it enables the kind of large-scale systematic comparison presented here, which would be difficult when methods are scattered across different toolboxes with incompatible interfaces and data formats. The bundling of simulation, solving, and evaluation into one package ensures that all methods are compared under identical conditions—same forward model, same noise realizations, same metrics—eliminating confounds that arise from cross-toolbox comparisons.

### 8.3 Design Decisions and Limitations

All solvers in `invertmeeg` are re-implementations based on published algorithms, not wrappers around original authors’ code. While we have validated the implementations against published results where possible, subtle differences may exist. Users who require exact reproduction of a specific paper’s results should verify against the original implementation.

The benchmark is intentionally narrow. It is EEG-only, uses synthetic simulations only, and fixes a single BioSemi-32 montage, a single `ico3` source space (1,284 dipoles), and a fixed-orientation forward model. These results should therefore be interpreted as valid within this benchmark family rather than as universal rankings for all EEG inverse problems.

The four scenarios span more than one source model, but they are not assumption-free. The focal and multi-source conditions use diffusion-basis patches with different order ranges, whereas the extended-source condition uses contiguous Gaussian patches; the low-SNR condition is defined by its noise regime rather than by a separate spatial model in Figure 3. This reduces but does not eliminate the possibility that some solver families are advantaged by similarity between their internal priors and the simulation family.

The novel solvers were developed under the same broad simulation assumptions used in the benchmark, even though the evaluation instances themselves were regenerated with different random draws. This means the frozen snapshot is informative about within-family generalization, but external validity to different EEG montages, source spaces, or real data remains limited.

The package currently supports only fixed-orientation source models. Free-orientation forward models are automatically converted to fixed orientation, which may not be appropriate for all applications.

Computational cost varies substantially across solver categories. Linear solvers (MNE, eLORETA) compute an explicit inverse operator and are fast to apply; iterative solvers (Champagne, BCS) may require many iterations to converge; and deep learning solvers require training, which can take minutes to hours. The neural-network baselines reported here were trained with reduced model capacity and training budget to keep the benchmark tractable; stronger ANN performance may be achievable with larger models and longer training, but that was not evaluated in this frozen snapshot. `invertmeeg` does not currently provide GPU acceleration for non-neural-network solvers.

The package is licensed under the GPL-3.0-only license.

### 8.4 Future Work

Planned extensions include GPU-accelerated linear algebra for large-scale source spaces, time-frequency source imaging methods, connectivity-informed inverse solutions, and integration with parcellation-based analyses. We plan to extend the benchmark to MEG sensor configurations and higher-density EEG montages, and to include real-data validation on established phantom and auditory/visual paradigm datasets.

We also plan to analyze the role of AI coding agents in solver prototyping more systematically. Several agent-assisted solvers rank strongly within this benchmark, but that observation should be interpreted cautiously until it is tested under held-out simulation families and real-data validation. A companion paper dedicated to that development workflow is forthcoming.

## 9 Conclusion

We have presented a frozen EEG benchmark of 106 inverse solvers across 4 scenarios, together with `invertmeeg`, a unified Python package that currently exposes 118 solver implementations through the `MNE-Python` ecosystem. Within this benchmark family, solver choice has a large effect on reconstruction quality, and the best-performing methods depend on source geometry, source multiplicity, and noise level.

Key findings include: (1) flexible subspace scanners and hybrid switching methods occupy the top of the frozen global ranking; (2) Bayesian refinements such as Subspace-SBL remain especially competitive in the noisy and extended-source regimes; (3) classical minimum norm defaults rank below several alternatives under this specific EEG setup; and (4) the benchmarked deep-learning baselines remain well behind the strongest classical solvers under the constrained training budget used here.

Not every package solver is benchmarked in this paper, but all 118 registered implementations are available in `invertmeeg`, together with the simulation framework and evaluation tools needed to reproduce and extend the EEG benchmark. The source code is available at https://github.com/LukeTheHecker/invertmeeg under the GPL-3.0-only license and can be installed via `pip install invertmeeg`. Documentation and tutorials are available at https://lukethehecker.github.io/invertmeeg/.

## A Detailed Benchmark Results

The following tables provide per-metric results for each of the four evaluation scenarios. Values are reported as means across 50 independent simulations per scenario, sorted by EMD. Metrics: EMD (earth mover’s distance in mm, primary), AP (average precision, secondary), MLE (mean localization error in mm), Corr (correlation).

## B Complete Solver Reference

The complete package registry is summarized below and provided as a machine-readable CSV in supplement/solver_reference.csv.

## Notes

### Competing Interest Statement

The authors have declared no competing interest.

### Summary of Updates

Clarified evaluation metrics, moved tables to appendix, improved figures.

